# Spatiotemporal transcriptomic map of ischemic brain injury

**DOI:** 10.1101/2023.03.28.534553

**Authors:** Daniel Zucha, Pavel Abaffy, Denisa Kirdajova, Daniel Jirak, Miroslava Anderova, Mikael Kubista, Lukas Valihrach

**Affiliations:** Laboratory of Gene Expression, Institute of Biotechnology of the Czech Academy of Sciences, Vestec, Czech Republic; Department of Informatics and Chemistry, Faculty of Chemical Technology, University of Chemistry and Technology, Prague, Czech Republic; Department of Cellular Neurophysiology, Institute of Experimental Medicine of the Czech Academy of Sciences, Prague, Czech Republic; Department of Radiodiagnostic and Interventional Radiology, Institute of Clinical and Experimental Medicine, Prague, Czech Republic; Faculty of Health Studies, Technical University of Liberec, Liberec, Czech Republic; TATAA Biocenter AB, Gothenburg, Sweden

## Abstract

The role of non-neuronal cells in the resolution of cerebral ischemia remains to be fully understood. To decode key cellular processes that occur after ischemia, we performed spatial and single-cell transcriptomic profiling of mouse brain tissue during the first week of injury. Cortical gene expression was severely disrupted, being defined by inflammation and cell death in the lesion core, and glial scar formation on the periphery. For each of the three major glial populations, an inflammatory-responsive state, resembling the reactive states observed in neurodegenerative contexts, was documented. The recovered spectrum of ischemia-induced oligodendrocyte states supports the emerging hypothesis that oligodendrocytes actively respond to and modulate the neuroinflammatory stimulus. Thus, we present a landmark transcriptomic dataset that provides a comprehensive view of spatiotemporal organization of processes in the post-ischemic brain and documents the conservation of glial response in CNS pathology.

## Introduction

Ischemic stroke is an acute pathological condition of a critical reduction in blood flow, caused by either sudden or gradual occlusion of cerebral arteries. Yearly it affects over 12 million people world-wide, with approximately 6.5 million deadly cases, making it the second most prevalent cause of death^1^. Long-term patient disability, and related medical and rehabilitation costs are additional major societal and financial burdens^2,3^. Although this creates a demand on development of new treatment strategies^4^, early mechanical thrombectomy and thrombolysis remain the sole approved therapies, applicable only in a short therapeutic window lasting several hours after the injury^5^.

Ischemic stroke is a complex injury, characterized by a cascade of pathological events involving interactions between a large number of cell types^6,7^. The dominant pathophysiological process is represented by death of neurons, accompanied by extensive inflammatory response. The inflammation is mediated predominantly by blood-borne immune cells, which invade the lesion parenchyma, joined by activated brain-resident cells^8^. The activation of resident glial cells leads to adoption of a phenotype spectrum, exhibiting a range of pro-inflammatory or anti-inflammatory properties^9^. The beneficial or detrimental role of both is largely determined by temporal and spatial factors as well as interaction with other cell types. This creates a unique opportunity for the latest omics technologies and advanced computational methods to provide holistic understanding of the post-ischemic molecular mechanisms.

Here, we characterized the acute phase of the experimental ischemic brain injury using the combination of spatial, single-cell and bulk transcriptomics to identify key transcriptional programs and cell types responding to the injury. We focused on ischemic lesion core and surrounding penumbra, where dramatic changes in cellular function and composition occur during the first days after the injury. As a dominant process, we further explored glial cell activation and compared it with the response in a neurodegenerative context. Our work represents an important point of reference for other studies as well as starting point in search for novel targets for stroke modulation in the period out of the current window for the therapy.

## Results

### Ischemic brain injury severely disrupts cortical gene expression landscape

To gain comprehensive view on genome-wide expression changes during progression of ischemic brain injury, we performed spatial transcriptomic analysis in 3-months old mice using an experimental model of permanent middle-cerebral artery occlusion (MCAO). The model mimics the majority of clinical stroke cases, which do not receive appropriate treatment in the narrow therapeutic window, leaving the clogged artery impassable^2,10,11^. We focused on the acute phase of the injury in a series of four timepoints: no injury (ctrl), and day 1, 3 and 7 post injury (DPI) (Fig. 1A).

**Figure 1.**
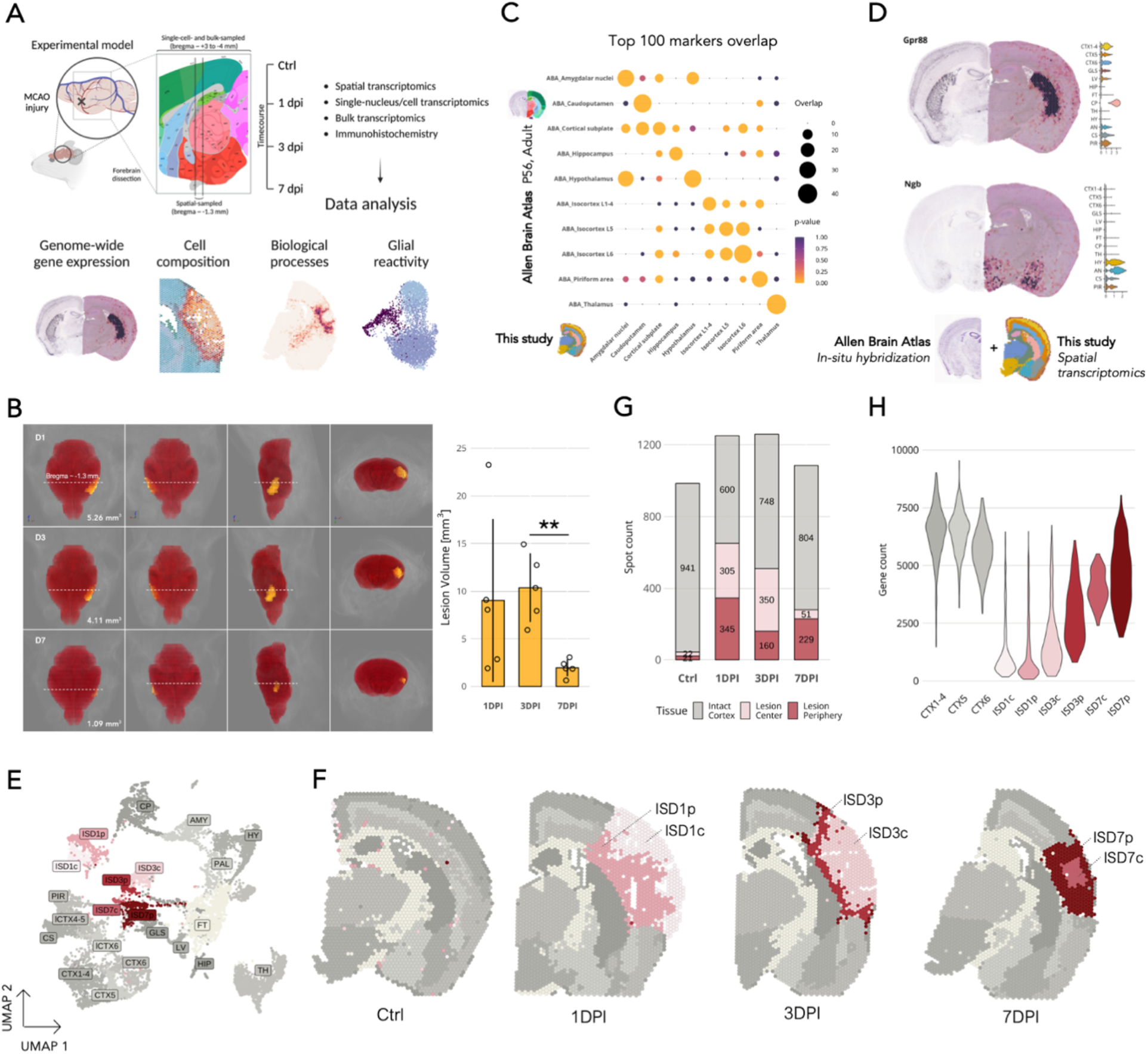
Cortical gene expression landscape undergoes major changes in the post-ischemic brain. A) Experimental design. B) Left: 3D visualization of repeated MRI scans of post-MCAO mouse brain at selected timepoints (rows) with the lesion highlighted in yellow. Lesion volume is recorded in the bottom-right corner of the first panel. The four panels in the left-to-right order show four views in the following orientations: top-down, bottom-up, side, and front view. An approximate location of the section collected for spatial transcriptomics, bregma -1.3 mm, is delineated with a dashed line. Bar plot on right: Mean volume of the post-MCAO lesions measured with MRI (n = 5 per timepoint). Error bars show standard deviation (st.dev). Wilcoxon’s rank sum test was used to assess statistical differences; the only significant difference is shown (3DPI vs., 7DPI, \*\**P* = 0.008). C) Overlap of the top 100 region-specific genes (fold-change sorted) identified in the ISH Allen Brain Atlas (Supp. Tab. 1). The size (circle size) and the significance (color scale, FDR-adjusted p-value) of the overlaps are indicated. D) Selected region-specific genes visualized in side-by-side comparison of the ISH Allen Brain Atlas image (left) with this study (right). Region-specific normalized expression is shown in the right-side violin plot. Additional side-by-side comparisons are provided in Supp. Fig. 3. E) UMAP of spots highlighting the brain region clustering. Ischemia-induced clusters are highlighted in shades of red. Abbreviations: amygdala, AMY; caudoputamen, CP; cortical subplate, CS; cortex layers 1-4, CTX1-4; cortical layer 5, CTX5; lateral cortex layers 4-5, lCTX4-5; cortical layer 6, CTX6; lateral cortex layer 6, lCTX6; fiber tracts, FT; glia limitans superficialis, GLS; hippocampus, HIP; hypothalamus, HY; ischemic region day 1 center, ISD1c; ischemic region day 1 periphery, ISD1p; ischemic region day 3 center, ISD3c; ischemic region day 3 periphery, ISD3p; ischemic region day 7 center, ISD7c; ischemic region day 7 periphery, ISD7p; lateral ventricle, LV; pallidum, PAL; piriform area, PIR; thalamus, TH. F) Spatial plot of the UMAP clusters from the panel E. Abbreviations: control, Ctrl; 1 day post injury, 1DPI; 3 days post injury, 3DPI; 7 days post injury, 7DPI. G) Number of spots spanning the uninjured and the lesioned cortexes. H) Gene count in the cortical regions. Spatial plots are in Supp. Fig. 1.

To ensure collection of representative ischemic brain samples for spatial transcriptomics, we first performed a pilot experiment assessing variability and reproducibility of the injury introduced by MCAO. We screened five mice per time-point using magnetic resonance imaging (MRI) and quantified the size of the brain swelling (Fig. 1B). The lesion volume was the most variable on day 1, averaging at 9.03 mm^3^ (st.dev ± 8.55 mm^3^). On day 3, we observed maximal average lesion volume at 10.38 mm^3^ with considerably smaller variability (± 3.61 mm^3^). Lesions measured on day 7 significantly retracted in volume, averaging 1.95 mm^3^ in volume (± 0.88 mm^3^). One additional mouse underwent repeated MRI-screening, confirming the regression of lesion volume in time (Fig. 1B). Guided by these results, we collected representative coronal brain cryosections from a location bregma -1.3 mm ± 0.1 mm and processed them on the Visium platform with 55 µm spot resolution from 10X Genomics.

First, we set out to provide baseline characterization of the profiled brain regions in a control section. The control section spanned a total area of 2417 spots, with median of ∼25,000 transcripts and ∼6,000 genes captured per spot (Supp. Fig. 1). Using *Seurat* spatial transcriptomics pipeline^12^, we iteratively clustered its spots at varying resolutions, and mapped the resulting UMAP clusters onto spatial plot (Methods). We obtained an optimum with 13 clusters corresponding to the mouse brain anatomy referenced in Allen Brain Atlas^13^ (Supp. Fig. 2). Although more detailed clustering could be achieved reminiscent of more detailed atlases^14^, we preferred the broader annotation for easier interpretability. To identify genes driving the anatomical distinction, we started with the differentially expressed genes (DEGs) between the annotated regions. For most regions we obtained hundreds of DEGs with fold-change > 1.5 and Bonferonni-adjusted *P*_*adj*_ < 0.01, Supp. Tab.1, exceptions being the cortical subplate region (87 markers) and the brain’s surface glial layer (88 markers). Although most markers differed in the level of expression, for some we observed strong regional specificity (Supp. Fig. 2). Additionally, the results included markers of specific cell types, e.g., *Pvalb, Lamp5, Plp1*, and *Ttr*, reflecting the regional cell type composition rather than a difference in expression level^15^. To test the robustness of the discovered markers, we overlapped the top 100 markers we identified with those in the Allen Brain Atlas *in-situ hybridization* (ISH) dataset (Supp. Tab. 1), for which we observed significant overlaps across all the regions (Fig. 1C), highlighting the conservation of region identity at the anatomical and transcriptional level. Side-by-side comparison of selected markers with the *ISH* images in Allen Brain Atlas also confirmed their spatial agreement (Fig. 1D; Supp. Fig. 3).

**Figure 2.**
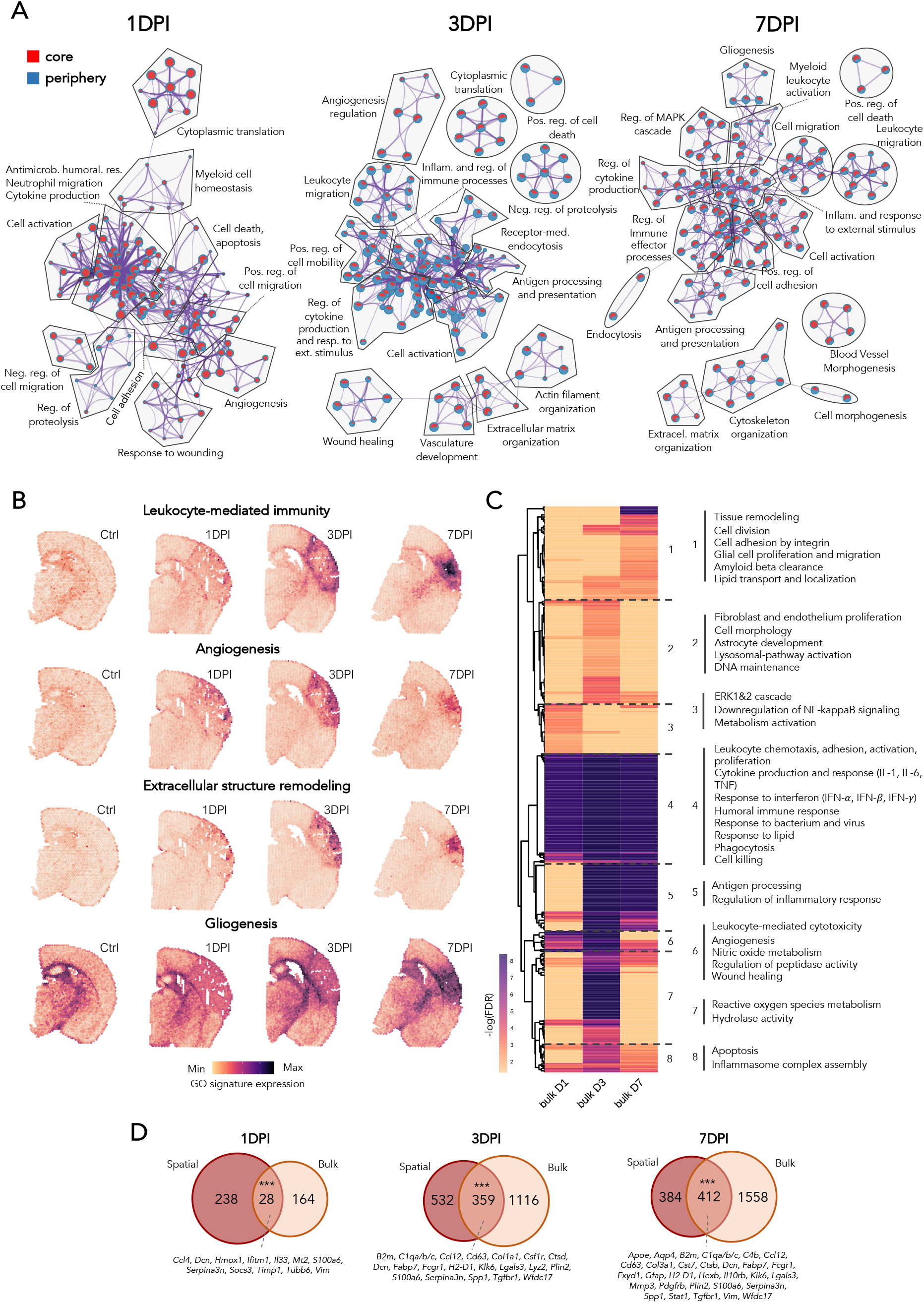
Acute phase of ischemic brain injury is characterized by persistent immune response and early activation of reparative processes. A) Gene ontology (GO) network of enriched biological processes based on the spatial transcriptomic data, grouped by the parent GO term (full list in Supp. Tab. 2). Each dot represents one enriched GO term, indicating the number of contributing DEGs (dot size) and core-periphery contribution (red-blue color scale, respectively). Prepared with Metascape^27^. B) Expression signatures of selected GO terms. C) Enriched GO biological processes from bulk RNA-seq. Selected representative GO terms of the grouping are highlighted on the side (full list in Supp. Tab. 3). D) Overlap of upregulated genes (DEGs with fold change > 1.5 and *P*_*adj*_ < 0.01) identified in the spatial dataset (union of core and periphery markers) and the bulk dataset, with selected overlapping genes listed. Significance of the overlap displayed as hypergeometric p-value (*\*\*\*P* < 0.001).

**Figure 3.**
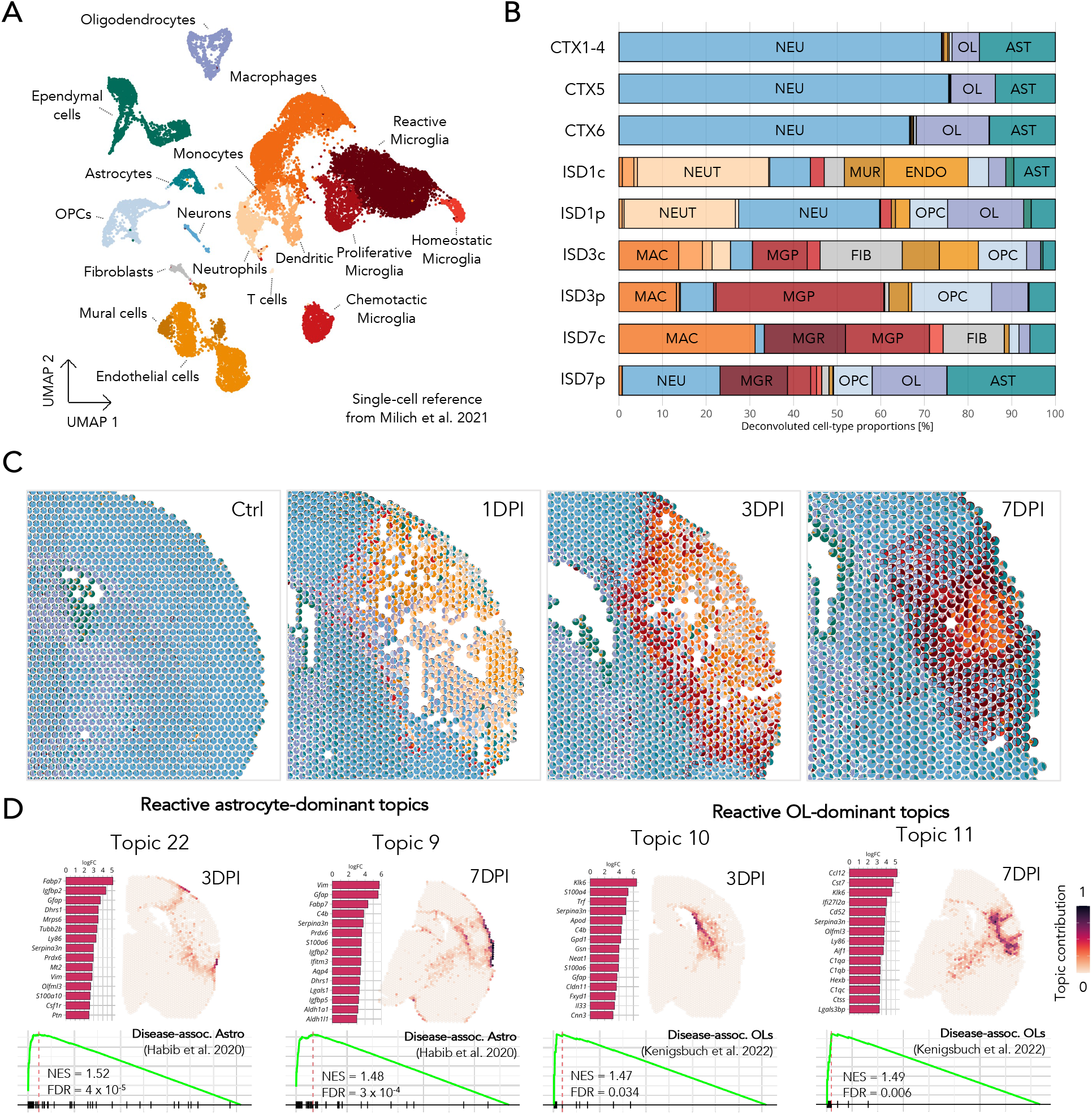
Deciphering cell type response to the ischemic injury. A) UMAP of cell populations from post-injury spinal cords, identified in the re-analyzed dataset of Milich et al.,^30^ with the same experimental design as in this study (n = 22170 cells). B) Average cell type proportion in the uninjured and ischemic regions as deconvoluted by RCTD^28^. Abbreviations: astrocytes, AST; endothelial cells, ENDO; fibroblasts, FIB; oligodendrocyte, OL; oligodendrocyte precursor cells, OPC; macrophages, MAC; microglia proliferative, MGP; microglia reactive, MGR; mural cells, MUR; neuron, NEU; neutrophils, NEUT. C) Detail view of the cortex with cell type proportions highlighted in the individual spots. The color labels correspond to panels A and B. D) Topics of expression programs identified by STdeconvolve^51^ with significantly enriched gene signatures of disease-associated astrocytes and oligodendrocytes (DAA^36^, DOLs^46^). Top topic marker genes are shown on the left-side, and enrichment plots with normalized enrichment score (NES) and FDR-adjusted p-value shown below.

Next, we integrated data from the control and ischemic sections to describe changes in the cortical gene expression introduced after the stroke. The profiled ischemic sections spanned a generally similar area of ∼2600 spots, with median of ∼ 12,000 transcripts and ∼4000 genes captured per spot (Supp. Fig. 1). Clustering revealed that the cortical lesion areas separated primarily based on time (ischemia day, ISD1, ISD3 and ISD7), and secondary based on the central (c) or peripheral (p) location within the lesion (Fig. 1E-F). The strict time-dependent clustering was interrupted by few tens of spots from ISD7p appearing on the very border of the day 3 lesion area, suggesting gradual resolving of the injury from the periphery to the center. Additionally, the ischemia-affected clusters spatially corresponded with brain areas supplied by middle cerebral artery^16^. Although the captured cortical area was approximately similarly sized with ∼ 1000 spots in each of the sections, the timepoints varied greatly in the area of lesioned parenchyma. On day 1, the lesion covered 52 % of the total cortical area, which declined to 40 % on day 3, and ultimately to 25 % on day 7 (Fig. 1G). This agrees with the pilot MRI screens and highlights the correlation between two independent technologies. The dynamic nature of the ongoing processes was also reflected in the overall transcriptional activity, expressed as detected genes per spot (Fig. 1H, Supp. Fig. 1). While the lesion on day 1 showed the lowest number of expressed genes, the activity increased with time and peaked on day 7 but did not reach the signal of uninjured cortex. This indicates an incomplete recovery and/or changes to transcriptional programs and cell composition. Interestingly, we also observed differences in the overall transcriptional activity between the central and the peripheral areas, supporting our previous notion on gradual resolution of the injury from periphery to center^6,11,17,18^.

The initial analysis of the control brain section demonstrated good quality of the collected data and anatomical agreement with the reference atlases. Analysis of the ischemic sections revealed severe disruption of the cortical transcriptional profile, with distinctive lesion core and enveloping penumbra.

### Inflammation and tissue remodeling are hallmarks of the early post-ischemic phase

To provide functional characterization of the ischemic injury in time and space, we calculated DEGs for each lesion cluster (Supp. Tab. 2), performed enrichment analysis (Supp. Tab. 2) and visualized the upregulated biological processes for each time point in a separate network, highlighting contributions of peripheral and central lesion areas (Fig. 2A). The most prominent clusters of the day 1 network indicated the upregulation of processes associated with leukocyte migration, adhesion, cytokine production, cell activation, and response to wounding, altogether reflecting increased infiltration and activation of peripheral immune cells^6,8,19,20^ (Fig. 2B). Concomitantly, terms related to cell death, apoptosis, proteolysis, extracellular matrix organization and angiogenesis suggested early initiation of reparative processes^21–23^ (Fig. 2B). Most processes were primarily activated in the central lesion, which likely reflects milder or delayed damage of the peripheral area^11^. This is consistent with the gradient-like character of the ISD1 peripheral cluster, which contrasts the enveloping-like shape observed at later time points (Fig. 1F). Many of the processes initiated on day 1 were still active on day 3, frequently even more (Fig. 2A, Supp. Tab. 2). These include persistent inflammatory response and processes restoring the integrity of the damaged tissue. However, in contrast to the previous time point, these two broad classes of processes showed distinct spatial localization. While the reparative machinery was more prominent in the central area, inflammation took place primarily in the periphery and involved immune cell activation and glial cell migration^9,17^ (Fig. 2B). A striking change was seen on day 7, when most processes, including inflammation, were localized to the central lesion (Fig. 3A). In the periphery, gliogenesis, axon ensheathment and synapse pruning were more pronounced, likely being first signs of neural reorganization^18,24,25^. Notably, while gliogenesis occurred in both lesion areas, detailed analysis revealed enhanced microglia activation in the center and astrocyte differentiation in the periphery, reflecting their respective localization (Supp. Fig. 4, Supp. Tab. 2). Interestingly, the peripheral and central lesion areas on day 7 showed a distinct difference in the number of DEGs (401 vs 1,905; *P*_*adj*_ < 0.01, fold change < -1.5 for downregulated and fold change > 1.5 for upregulated genes; Supp. Fig. 4, Supp. Tab. 2), confirming profound transcriptional changes in the lesion core and stabilization of the injury in the periphery.

**Figure 4.**
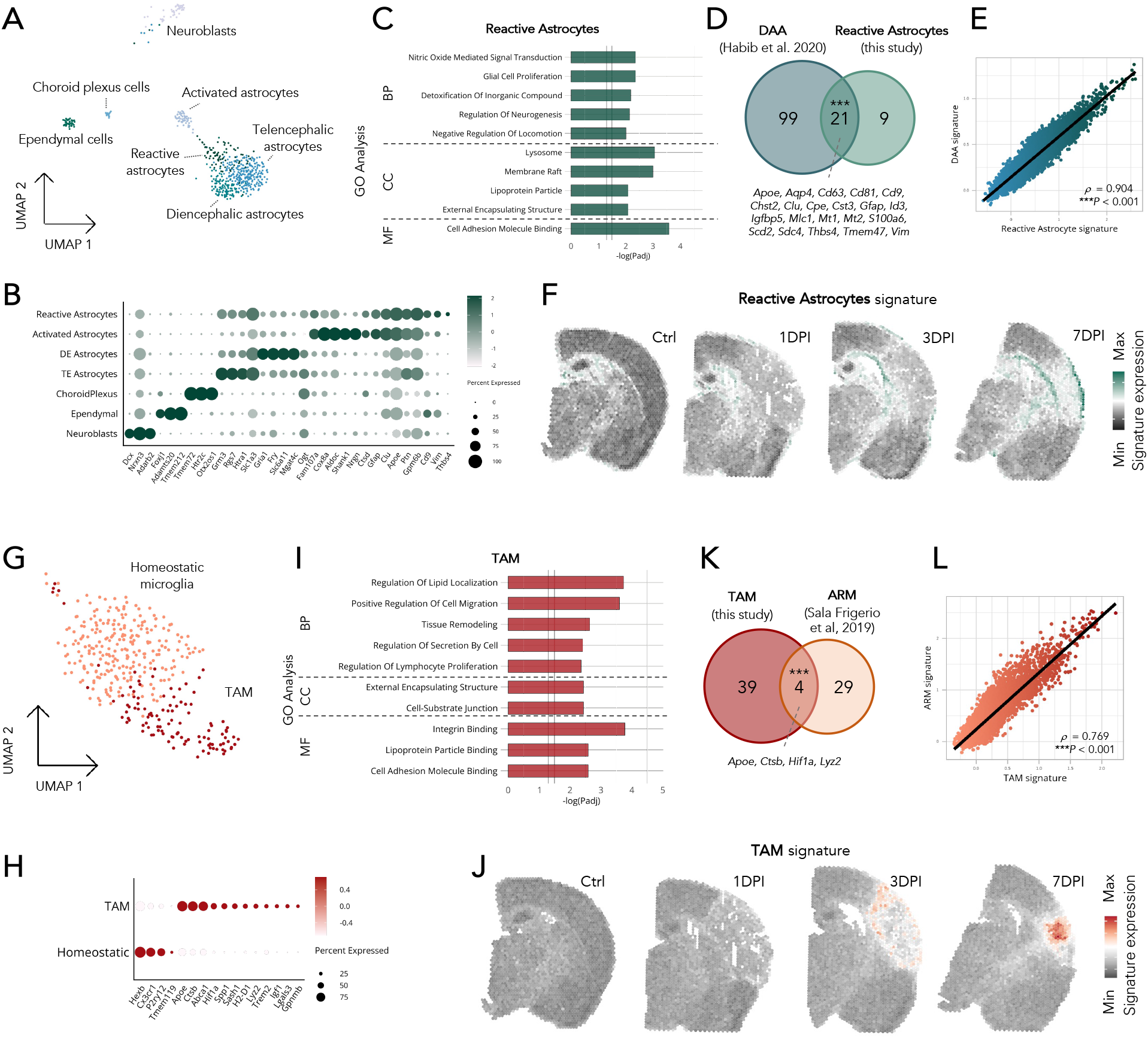
Astrocytes and microglia acquire reactive states in response to the post-MCAO conditions. A) UMAP of astroependymal populations from post-MCAO mouse brains (n = 610 nuclei). B) Marker genes of astroependymal populations. Expression level (color scale) and the percentage of population expressing the marker (dot size) are shown. C) Representative enriched gene ontology (GO) terms from the reactive astrocytes markers, categorized as ontology terms belonging to biological processes (BP), cellular component (CC) or molecular function (MF). FDR-adjusted *P* value is shown. D) Shared markers between the signatures of disease-associated astrocytes (DAA)^36^ and this study’s reactive astrocytes. Hypergeometric p-value *\*\*\*P* < 0.001. E) Correlation between the the expression signatures of this study’s reactive astrocytes (x axis) and the DAA signature (y axis) in the spatial dataset, quantified with quantified with Spearman’s rank correlation coefficient *ρ*. F) Spatial reactive astrocytes signature is shown, calculated using the top 15 population marker genes (fold-change sorted) quantified as the difference between the average expression of this signature and a signature of iteratively randomly selected genes of similar expression levels (Methods). G) UMAP of microglial populations from post-MCAO mouse brains (n = 362 nuclei). H) Marker genes of microglial populations. Expression level (color scale) and the percentage of populations expressing the marker (dot size) are shown. I) Representative enriched gene ontology (GO) terms from the reactive astrocytes markers, categorized as ontology terms belonging to biological processes (BP), cellular component (CC) and molecular function (MF). FDR-adjusted *P* value is shown. J) Spatial trauma-associated microglia (TAM) signature is shown, calculated using the top 15 population marker genes (fold-change sorted) iteratively quantified as the difference between the average expression of this signature and a signature of randomly selected genes of similar expression levels (Methods). K) Shared markers between the signatures of this study’s TAMs and activated response microglia (ARM)^45^, hypergeometric p-value *\*\*\*P* < 0.001. L) Correlation between the expression signatures of this study’s TAMs (x axis) and the ARM signature (y axis) in the spatial dataset, quantified with Spearman’s rank correlation coefficient ρ.

To validate the spatial transcriptomic measurements, we collected lesion areas from a minimum of four mice per time point and performed genome-wide transcriptional analysis using bulk RNA-sequencing (RNA-seq). Also for the bulk data we calculated DEGs and performed functional characterization by gene-set enrichment analysis (GSEA)^26^ (Fig. 2C, Supp. Tab. 3). The GO terms enriched across time points were associated with cell activation and immune response (e.g. leukocyte migration, cytokine production, cell-cell adhesion), largely mirroring observations in the spatial data. In contrast, processes related to reparative processes (e.g. extracellular matrix organization and angiogenesis, Fig. 2B), which were detected already on day 1 in the spatial data, were found upregulated not until day 3 in the bulk experiment. This is likely due to the rather small contribution from the central lesion area, which is the initiation site of the reparative processes, to the bulk dataset and therefore remains unnoticed. To obtain global view on the concordance of the spatial and bulk measurements, we merged central and peripheral areas for each time point in the spatial data set, combined their DEGs and compared with the DEGs from the bulk RNA-seq (Fig. 2D, Supp. Tab. 2 and 3). Overlap at each time point (hypergeometric *\*\*\*P* < 0.001) reflects agreement of the measurements, thus confirming the spatial data.

Altogether, the functional characterization of the ischemic lesion revealed dramatic inflammatory response with many components and distinct spatial localization over time. Reparative activity was found predominantly in the center lesion, while early signs of neural reorganization were seen in the periphery one week after the injury. Processes and their temporal dynamics measured in bulk RNA-seq supported the observatsions made using spatial transcriptomics.

### Deconvolution revealed dramatic changes in cell type composition in the ischemic lesion

Changes introduced by ischemic brain injury are orchestrated by multiple cell types^6^. To complement the previous analysis with cell-type specific context, we applied computational deconvolution and assessed changes in relative proportion of cell types in the injured cortex. We applied high-ranked RCTD algorithm^28,29^, which uses single-cell reference to estimate cell-type content in the individual spots of spatial assays. As the quality of the reference determines the quality of the deconvolution, we chose the recent atlas of mouse spinal cord injury^30^, for its similar cell type composition and experimental settings. The re-analysis of the reference dataset gave 17 cell populations (Fig. 3A, *Methods*), which were annotated based on the expression of canonical markers (Supp. Fig. 5). The matrix was then used to deconvolute the data to obtain cell type contents in the individual sections.

**Figure 5.**
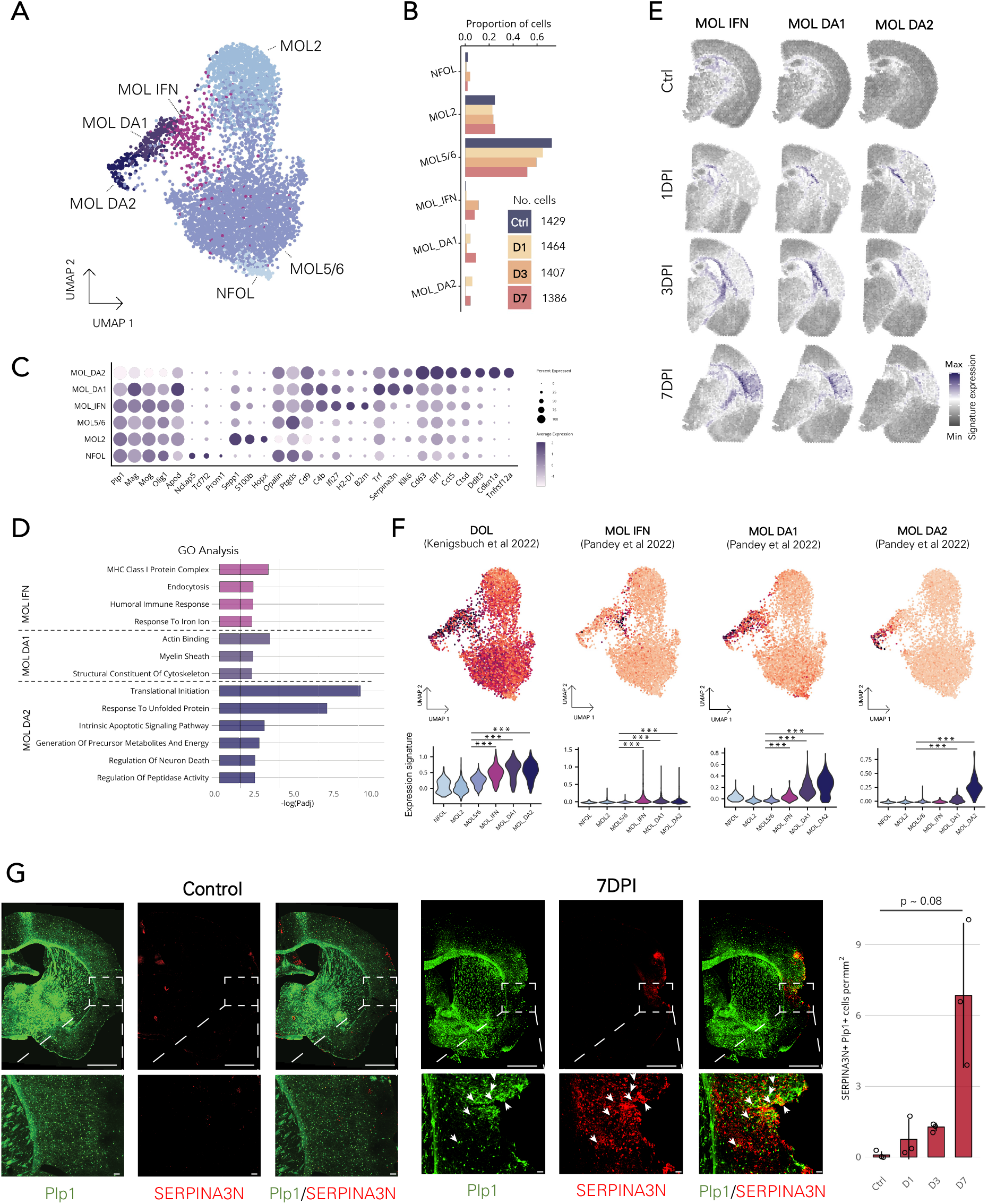
Ischemic conditions induce a spectrum of reactive states in oligodendrocytes. A) UMAP of populations in the integrated dataset of single nuclei (n = 1113) and single cells (n = 4673) of post-MCAO oligodendrocytes. Abbreviations: Newly formed oligodendrocytes, NFOL; mature oligodendrocytes type 2, MOL2; mature oligodendrocyte types 5 and 6, MOL5/6; interferon-responsive mature oligodendrocytes, MOL IFN; disease-associated mature oligodendrocytes 1, MOL DA1; disease-associated mature oligodendrocytes 2, MOL DA2. B) Proportions of oligodendrocyte populations at the respective timepoints. C) Marker genes of oligodendrocyte populations. Expression level (color scale) and the percentage of populations expressing the marker (dot size) are shown. D) Representative enriched gene ontology (GO) terms for trauma-associated oligodendrocytes, categorized as ontology terms belonging to biological processes (BP), cellular component (CC) or molecular function (MF). FDR-adjusted *P* value is shown. D) Spatial trauma-associated oligodendrocyte signatures are shown, calculated using the top 15 population marker genes (fold-change sorted), quantified as difference in the average expression of this signature and a signature of iteratively randomly selected genes with similar expression levels (Methods). F) Expression signatures of disease-associated populations (color scale) identified in the studies of Kenigsbuch et al.,^46^ and Pandey et al.,^34^ that characterized oligodendrocyte states in models of Alzheimer’s disease and multiple sclerosis. Concrete values are shown in the violin plots below, with the significance of differences assessed using Wilcoxon rank sum test with FDR-adjusted *P* values (*\*\*\*P* < 0.001). G) Coronal mouse brain sections showing mature oligodendrocytes (*Plp1+*) and serine-protease inhibitor (SERPINA3N+); a marker of neuroinflammatory response. Average cell count in three sections per mice, and three mice per condition. Wilcoxon rank sum test was used for statistical testing (n = 3), error bars indicate st.dev. Scale bars: 1000 µm and 50 µm for the hemisphere and the inset image, respectively.

The analysis uncovered dramatic changes in the cell type composition (Fig. 3B, Supp. Fig. 5). While the uninjured cortex was composed mainly of neurons, astrocytes and oligodendrocytes, the injury caused a substantial decline in neurons, triggered infiltration of peripheral immune cells, and proliferation of glia, vascular cells and fibroblasts. Most changes were coordinated in time and space. Neuronal death occurred on day 1 in a gradient from center to the periphery, suggesting gradual expansion of the injury^11^. It peaked on day 3, with a minor difference between the central and peripheral lesion. This changed on day 7, when the neuron count increased in the periphery, indicating initiation of neural reorganization. This is in line with the previous spatial and bulk RNA-seq results (Fig. 2A and C, Supp. Tab. 2), as well as the reduction of the lesion size quantified by MRI (Fig. 1B). While peripheral immune cells were not detected in the uninjured cortex, they dominated the lesion on day 1, with largest contribution being neutrophils. Other peripheral immune cells (monocytes, dendritic cells, macrophages, T cells) were less abundant and showed sparse localization within the lesion (Supp. Fig. 5). The neutrophils were replaced by macrophages on day 3, which further expanded to the central lesion on day 7, while almost disappeared from the periphery. The immunological niche dominating the lesion center on day 3 also contained monocytes, neutrophils, and dendritic cells, which were largely missing from the periphery at this time point. This strong immune response was also seen in the gene ontology analysis of both spatial and bulk RNA-seq data (Fig. 2A and C, Supp. Tab. 2). Vascular cells and fibroblasts, reflecting angiogenesis and the formation of the fibrotic scar, also displayed distinct time and spatial profile. Whereas angiogenesis was developed already on day 1 in the central lesion, fibrosis developed on day 3 and was restricted to the central lesion area.

Glial cell proliferation and differentiation were other prominent processes that the deconvolution analysis captured (Fig. 4B-C, Supp. Fig. 5). Focusing on the three major glial populations: astrocytes, oligodendrocytes and microglia, we observed significant decrease in the fraction of astrocytes and oligodendrocytes on days 1 and 3, which largely reversed in the periphery on day 7, reflecting the formation of the glial scar surrounding the central lesion^17,31,32^. Gradual decrease in the oligodendrocyte fraction was counteracted by rise of oligodendrocyte precursors (OPCs), which were abundant in the peripheral lesion on day 3 and 7, and largely disappeared in the central lesion on day 7. Their differentiation potential is, however, not restricted only towards oligodendrocytes; OPCs are also known to produce astrocytes after injury^17,33^. The most striking temporal changes were observed for microglia. These CNS resident immune cells were hard to localize in the uninjured cortex by deconvolution. In the lesion they were easily identified and showed subtle grow on day 1, followed by steep increase on days 3 and 7, when they became the most abundant cell type. The observed trends were confirmed by visualizing expression intensity of canonical cell type-specific genes (Supp. Fig. 4), as well as by immunostaining, measuring the signal of GFAP^+^ in astrocytes, PGDFRα^+^ in OPCs, PLP1^+^ in oligodendrocytes and IBA1^+^ in microglia (Supp. Fig. 6). We also visualized expression of selected markers of reactive glia identified in a range of neuropathologies^34–46^, e.g. *Ccl12, Spp1, Cstd, Cst7, Trem2, Apoe, Vim, Mt1, Serpina3n, C4b, Klk6*, and found them in a distinct spatial pattern inside and near the ischemic lesion (Supp. Fig. 4).

To sum up, the deconvolution analysis revealed dynamic changes in cell type distributions within the ischemic lesion. The responses organized in distinct spatiotemporal patterns, with an early contribution from the peripheral immune and vascular cells that is taken over in the acute phase by glial cells and fibroblasts.

### Ischemic brain injury triggers spatially localized activation of glial cells

As we found distinct spatiotemporal patterns for several genes associated with reactive populations of glial cells (Supp. Fig. 4), we focused further on their characterization. First, we estimated the relative changes of the four microglia populations: homeostatic, chemotactic, proliferative, and reactive, that were in the reference single-cell matrix used for the deconvolution (Fig. 3A-C, Supp. Fig. 5). Homeostatic microglia (*P2ry12, Hexb*) were hardly detected in any section, possibly due to the dominance of the other microglial populations. The few microglia on day 1 were chemotactic (*Ccl12, Ccl7*) and surrounded the central lesion. They are known to contribute to the infiltration and activation of peripheral immune cells as well as other glial cells^47–50^. Day 3 microglia were represented by a proliferative population (*Ube2c, Cenpa*), with highest abundance in the periphery that enclosed the lesion. The coordination of microglial activation and proliferation are hallmarks of microgliosis, which induces glial scar formation^48^. On day 7, the proliferative population was partially replaced by reactive microglia, characterized by upregulation of inflammatory signaling molecules (*C1qa/b/c, Ccl3, Ccl4*), lipid metabolism genes (*Apoe, Apoc1*), and lysosome-related genes (*Lyz2*) including cathepsins (*Ctsd, Ctsz*) and their inhibitors cystatins (*Cst7, Cst3*). Notably, many of the reactive genes were increasingly expressed in these microglial populations, indicating their gradual activation^37^.

To map the activation of astrocytes and oligodendrocytes, we used the reference-free deconvolution tool STdeconvolve^51^, which can identify topics of cell-type transcriptional profiles, estimates their contributions in each spot, and provides a list of the topic’s marker genes. In total, we defined 23 topics in the control section, 24 topics in the day 1 and day 3, and 21 topics in the day 7 section (Supp. Fig. 7), each characterized by a list of marker genes (Supp. Tab. 5). Many of the topics were representative of cell types and agreed with the reference-based deconvolution results (Supp. Fig. 5). Next, we screened the lists of topic marker genes for enrichment of signature genes of disease-associated astrocytes (DAA)^36^ and disease-associated oligodendrocytes (DOL)^46^, representing activated populations previously described in neurodegenerative contexts. Four topics localizing around the lesion showed significant positive enrichment (Fig. 3D). Topics 10 and 22 on day 3 and topic 9 on day 7 were enriched for both the DAA and DOL signatures, while topic 11 on day 7 was enriched only for the DOL signature. The overlapping enrichments of DAA and DOL, for example seen by *Serpina3n* and *C4b*, were expected, as the similarity of these reactive states has been reported in several neurodegenerative models^34,46^. These topics were also characterized, albeit to a lesser extent, by multiple microglial markers (*Hexb, Csf1r;* Supp. Tab. 5), illustrating their mutual relatedness. Interestingly, the reactive topics enveloped the lesion on day 3, but extended outside the glial scar to the white matter tracts on day 7. In concordance, myelin, a building compound of white matter tracts, is highly susceptive to damage upon sustained inflammation^52^ and glial cells are the key influencers of myelin health irrespective of pathological conditions^43,53–55^. Lastly, we visualized gene signatures of reactive microglia, astrocytes and oligodendrocytes from different pathological conditions, including Alzheimer disease (AD)^36,37,44–46^, demyelination^41,42^, aging^43^ and stroke,^40^ finding them localized to the area forming the glial scar (Supp. Fig. 8, Supp. Tab. 5).

Altogether, this analysis confirms the gradual activation of glial cells that leads to the initiation of the glial scar enclosing the ischemic core. The overlapping signatures of reactive glia suggest their close interaction as well as similarities in their expression programs.

### Single-nucleus profiling of post-ischemic brain reveals rise of reactive glia

The previous analyses gave strong evidence for the presence of reactive glial cells in the ischemic brain. To confirm, we performed single-nucleus RNA-seq (snRNA-seq) of the control and post-ischemic brains on days 1, 3 and 7. We sampled tissue in positions +3 to -4 mm to bregma to include comparable representations of brain regions to those analyzed above using spatial transcriptomics (Fig. 1A). In total, we obtained 7946 high-quality nuclei, which were classified into 11 major cell type populations, including clusters of glutamatergic and GABAergic neurons, neuroblasts and clusters of glial, ependymal, and vascular cells (Supp. Fig. 9). We further focused on three glial populations: 1) astroependymal cells (n = 763 nuclei), 2) microglia (n = 426 nuclei), and 3) oligodendrocyte lineage cells (n = 1433), which we analyzed in detail.

In the astroependymal cluster, we identified eight populations: neuroblasts (*Dcx*), choroid plexus cells (*Tmem72*), ependymal cells (*Foxj1*) and four astrocyte populations (Fig. 4A-B, Supp. Tab. 6). In concordance with Zeisel et al.^56^, two of the astrocytes populations represented homeostatic states, with region distinctive origin in telencephalon and diencephalon (Supp. Fig. 9). Another astrocyte population, named activated astrocytes, was characterized by high levels of ribosomal and mitochondrial reads, had active energy metabolism (*Cox8a, Aldoc*), and expressed genes involved in neuron maintenance and survival (*Shank1, Nrgn*; Supp. Tab. 6). These markers were predominantly enriched in hippocampal dentate gyrus, caudoputamen and subpial regions, suggesting possible neurogenic potential as described by Zamboni et al.,^57^ and Batiuk et al.^58^ (Supp. Fig. 9). The fourth astrocyte population showed reactive status, reflected by elevated expression of *Gfap, Apoe*, and *Clu*, and overexpression of genes related to cell adhesion (*Cd9, Ptn*), proliferation (*Vim, Thbs4*), lysosomal activity (*Ctsd*), and membrane raft lipids (*Gpm6b*) (Fig. 4C, Supp. Tab. 6). Their reactive profile was also supported by a significant overlap of 21 markers with Habib’s DAA population^36^ (hypergeometric *\*\*\*P* < 0.001, Fig. 4D), and correlation with the DAA signature in the spatial dataset (Spearman’s *ρ* = 0.904, *\*\*\*P* < 0.001 Fig. 4E). The reactive astrocytes showed telencephalic origin consistent with the cortical localization of the injury (Supp. Fig. 9).

Microglia separated into populations of homeostatic (*P2ry12)* and trauma-associated microglia (*Apoe, Spp1*; TAMs) (Fig. 4G). TAM-specific expression reflected their reactive profile (Fig. 4H), involving genes related to cell migration, proliferation, and adhesion (*Gpnmb, Sash1, Lgals3*), protein secretion (*Spp1, Igf1*), tissue remodeling (*Hif1a*), and lipid transport (*Abca1*, Fig. 4I, Supp. Tab. 6). TAMs were primarily enriched in the ischemic samples, peaking day 3 and comprising almost half of the collected microglia (Supp. Fig. 9). Spatial data confirmed the increase of the TAM signature in the lesion boundary on day 3 and its accumulation in the center on day 7 (Fig. 4J). As for the astrocytes, we investigated the similarity of TAM with the activated response microglia (ARM) described by Sala Frigerio^45^ in the 5XFAD mouse model of Alzheimer’s disease. The significant overlap of TAM and ARM signatures with 4 shared genes (*Apoe, Ctsb, Hif1a, Lyz2;* \*\*\**P* < 2 × 10^−6^, Fig. 4K) and the strong correlation of their signatures in the spatial datasets (Spearman’s *ρ* = 0.769, *\*\*\*P* < 0.001, Fig. 4L) suggested activation of similar transcriptional programs in acute and neurodegenerative pathology. Although we did not find the activated cluster of interferon-responsive microglia (IRM) described by Sala Frigerio in our snRNA-seq, we detected the distinct IRM signature in the central lesion on day 7 in our spatial datasets (Supp. Fig. 9).

Oligodendrocyte lineage cells were annotated using nomenclature of Zeisel et al.^56^ as reference. This yielded a total of four populations: oligodendrocyte precursor cells (OPC; *Pdgfra, Sox6*), newly formed oligodendrocytes (NFOL; *Tcf7l2, Prom1, Nckap5*), major population of mature oligodendrocytes (MOL; *Plp1, Grm3, Mast4*), and a MOL population expressing reactivity markers (*Serpina3n, C4b*), which we named trauma-associated oligodendrocytes (TAOs) (Supp. Fig. 9). TAOs were found exclusively in the ischemic samples and had elevated levels of long-noncoding RNAs (*Neat1, Pvt1*), and genes related to phagocytosis (*C4b, Pros1, Dock1*), cell division (*Pkp4, Cdc14a*), and positive regulation of immune effector processes (*Il33, C4b, Gab2*). Interestingly, oligodendrocytes (OLs) with similar profile have been observed in neurodegenerative context^34,46^. In the spatial dataset, TAO signature was enriched in the oligodendrocyte-rich white matter tracts in all sections, but in the ischemic sections primarily on the lesion boundary, progressively enveloping the lesion (Supp. Fig. 9). This spatial pattern was similar to the DOL-enriched topics (Fig. 3D), cross validating the previous deconvolution analysis.

To sum up, snRNA-seq confirmed the presence of reactive glial cell populations in the post-ischemic brain tissue. The identified subtypes showed similarity with the activated populations characterized previously in mouse models of neurodegeneration, highlighting shared features of transcriptional programs acquired by glia in response to pathophysiological stimuli.

### Spectrum of reactive oligodendrocytes emerges after ischemic brain injury

Prompted by increasing number of evidence suggesting immunomodulatory role of OLs in neurodegeneration^34,35,41,43,44,46,55^, we opted to further characterize OLs in the context of acute trauma. Using the same experimental design, we FACS-enriched *Plp1+* cells from the control and ischemic brain and performed single-cell RNA-seq (scRNA-seq). After QC, we identified 4573 high-quality OLs, which we integrated with the single nucleus dataset (Fig. 5A, Supp. Fig. 9). The increased number of data points allowed us to distinguish two mature OL clusters, MOL2 (*S100b, Hopx*) and MOL5/6 (*Ptgds, Opalin*), improve annotation of the NFOL cluster (*Tcf7l2, Prom1*), and recover three pathology-enriched MOL populations (Fig. 5B-D). Using the annotation introduced by Pandey et al.,^34^ we termed them interferon-responsive and disease-associated mature OLs 1 and 2 (MOL IFN, MOL DA1 and MOL DA2).

The population of MOL IFN was characterized by terms related to immune response, particularly to interferons (*Ifi27, Ifi27l2a*) and antigen presentation by MHC class I complex (*H2-D1, B2m*). This population also upregulated genes related to uptake of extracellular debris (*C4b, Cd9, Trf*), possibly in an effort to restore tissue homeostasis. Their spatial expression signature progressively strengthened (Fig. 5E), starting near the lateral ventricle and expanding along the lesion periphery to the white matter of corpus callosum and deeper fiber tracts. On day 7, the signature encircled the lesion core, suggesting their crosstalk with other cells forming the glial scar. MOL DA1 had elevated ribosomal RNA content (Supp. Fig. 9) and showed the highest expression levels of reactive marker genes – *Serpina3n, C4b*, and *Klk6*^46^ (Fig. 5C). GO analysis indicated their role in myelination and cytoskeletal restructuring (*Arpc1b, Tubb3*). The additional investigation of markers pointed to regulation of extracellular serine proteases activity (*Serpina3n, Klk6*), cell adhesion (*Cd9, Cd63*), metal ion homeostasis (*S100a6, S100a16, Calm2, Trf*), and enhanced lipid metabolism (*Apod, Fabp5*). Spatially, MOL DA1 overlapped with MOL IFNs, but their expression signature was less prominent in the deeper white matter fiber tracts (Fig. 5E). MOL DA2 represented the most distinct population of pathology-enriched MOLs, having high ribosomal activity and upregulated genes associated with translation initiation (*Eif1, Eif4a1*) and protein-folding chaperons (*Cct2, Cct5*; Fig. 5D, Supp. Fig. 9). Their activated profile was also characterized by upregulation of metabolic processes (*Cox5a, Atp5a1*) and regulation of peptidase activity (*Serpina3n, Ctsb, Ctsd, Timp1*). However, they also expressed genes related to stress (*Fos, Jun*), cell-cycle regulation (*Cdkn1a*), and apoptosis (*Gadd45b, Tnfrsf12a, Ddit3*), indicating a struggle for survival. Another distinct feature was low expression of myelinating genes (*Plp1, Mog, Mag*), suggesting reduced myelination capacity. The spatial expression signature was the most prominent on day 1 in corpus callosum near the lateral ventricle, slowly decreasing in time and residing in the outermost OL layer around the lesion (Fig. 5E).

The marker genes and functional annotation of pathology-enriched populations showed substantial similarity to those described in a neurodegenerative context^34,46^. Indeed, all three reactive populations were enriched for the DOL signature^46^, which we used for the annotation of single-nucleus data (Fig. 5F). Moreover, we also noticed that MOL IFN, MOL DA1 and MOL DA2 showed significant similarity to the OL populations identified across several models of Alzheimer’s disease and multiple sclerosis by Pandey et al.^34^. To confirm the presence of reactive oligodendrocytes, we selected *Serpina3n* gene as a prominent reactive marker shared by all three populations and used immunohistochemistry to quantify PLP1+ SERPINA3N+ cells up to 7 days post ischemic injury (Fig. 5G). While we did not detect PLP1+ SERPINA3N+ cells in control brains, we observed it increases after injury, peaking on day 7. In accordance with the spatial transcriptomics results, the positive signal accumulated in the lesion periphery enveloping the lesion core (Supp. Fig. 5). Notably, SERPINA3N+ was expressed also by other cell types, possibly including astrocytes, neurons, and OPCs^30,35,59,60^.

To sum up, combining snRNA-seq and scRNA-seq we identified a spectrum of reactive oligodendrocytes populations specific to the post-ischemic brain conditions. While all populations expressed reactive marker genes like *Serpina3n*, reflecting the potential to reduce neuroinflammation^60,61^, they differed in the expression of immunogenic genes and metabolic activity. They also showed specific spatial-temporal organization and similarities to reactive oligodendrocytes observed in neurodegenerative diseases.

## Discussion

In this study, we mapped the transcriptional landscape during the acute phase of experimental ischemic brain injury, providing a holistic view on the response mediated by a complex network of cellular components. We showed that anatomical brain regions as well as lesion areas, divided into central and peripheral zones, are robustly captured by spatial transcriptomics (Fig. 1). Cell activation, inflammation and tissue remodeling were hallmarks of coordinated response to ischemia (Fig. 2). The cellular response is organized into layers, resulting in the formation of a fibrotic core and a glial scar composed of activated populations of glial cells (Fig. 3). Using single-cell and single-nucleus transcriptomics, we characterized the trauma-associated profiles of glial cells, showing similarities to reactive states described in neurodegenerative CNS disorders (Fig. 4-5).

Spatial transcriptomics can accurately recover brain’s anatomy, owning to the unique function and cellular composition for each of its parts^13^. After ischemic injury, the cortical structure and function is severely disrupted, altering a part of the lesioned cortex into ischemic core and enclosing penumbra, which later transforms into the glial scar^17^. The ischemic core is the most affected area, whilst the penumbra maintains some of its metabolic and polarization capacity^7,11,62^. Indeed, we characterized the lesion core as the epicenter of cellular death accompanied by infiltration of peripheral immune cells, upregulating genes linked to cell activation, cytokine production and antimicrobial humoral response (Fig. 1-2, Supp. Tab. 2). Early infiltration of immune cells was followed by fibroblasts and vasculature cells to restore microvasculature and regulate the infiltration of peripheral immune cells^6^ (Fig. 2-3). Inflammation in the ischemic core acts upon astrocytes, microglia, oligodendrocyte lineage cells, triggering them to acquire activated states, leading to the formation of a dense limiting border known as glial scar^9,63^.

During the seven-day post-injury period, the glial scar matured, evidenced by the change in its thickness and cellular composition (Fig. 1-3). On day 1, we observed the expression gradient of chemokine genes (*Ccl12, Ccl2, Ccl3, Ccl4;* Supp. Fig. 4, Supp. Tab. 2), delineating lesion border from the brain parenchyma. Using reference-guided deconvolution analysis, we uncovered a small reactive microglial population on the lesion border, marked by upregulation of chemokine genes (Fig. 3, Supp. Fig. 5). Chemotactic microglia can produce inflammatory signal soon after acute injury^47^ or near demyelinated lesions of slowly progressing pathology^38^, which initiates microglial activation cascade that densifies the nearby tissue^48^. Another role is to guide locomotion of multiple cell types, including peripheral immune cells^64^, neuronal progenitor cells,^49,50^ as well as oligodendrocyte-precursor cells^65^ (Supp. Fig. 5 and 6). With chemokines, we observed a gradient of reactive genes (*Serpina3n, Gfap, C4b, Mt1;* Supp. Fig. 3), suggesting early activation of other glial cell types. The expression of *Serpina3n* is particularly intriguing, considering its inhibitory activity of Granzyme B protease, a potent neuronal toxin expressed by cytotoxic CD8+ T cells^61^. Under pathological conditions, *Serpina3n* can be expressed by astrocytes^36,44,60^, oligodendrocytes^35,43,46^, OPCs^30^ and neurons^59,60^, suggesting a general defense mechanism against T-cell mediated cytotoxicity. In concordance, we also detected multiple cellular sources of SERPINA3N (Fig. 5).

On day 3, the lesion area acquired more layered structure composed of different cellular components (Fig. 3). The lesion core undergone intensive remodeling, coordinated by a spectrum of immune cells, vasculature cells, and fibroblasts. The lesion periphery, almost fully enclosing the lesion core, consisted predominantly of proliferating microglia and OPCs. Their proliferation and activation-initiated hypertrophy densify the tissue, isolate the inflamed ischemic core from brain intact parenchyma and initiate the formation of the glial scar^17,48^. A question is how beneficial the microglia activation is to the ischemia outcome^18^. In a simplified view, the activated microglia adopt either pro-inflammatory or anti-inflammatory state (M1/M2)^66^, but recent publications report it is more complex^36–38,41,43–45,54^. The small population of trauma-associated microglia (TAM), which we recovered with snRNA-seq (Fig. 4), was characterized by genes (*Apoe, B2m, Lyz2*) expressed in early activated microglia identified in experimental models of AD^37,45^. These are considered beneficial in the early stages of neurodegeneration but become detrimental as the pathology progresses and activated microglia mature into the chronic reactive state (*Apoe*^*Hi*^, *Trem2*^*Hi*^, *Cst7*^*Hi*^). Interestingly, we also observed a gradual increase of the expression of these genes in the spatial dataset, peaking day 7 (Supp. Fig. 4), suggesting a similar mechanism of activation in the ischemic brain injury. Deconvolution analysis confirmed this finding, indicating reactive microglia that express the late reactivity markers, are the most abundant cell population on day 7 (Fig. 3, Supp. Fig. 5). Their role in the acute phase of the ischemic injury remains to be investigated, but is likely related to the beneficial effect of a similar population recently found studying spinal cord injury^67^.

The pathological milieu and the recruiting microglia affect nearby astrocytes and oligodendrocytes, which undergo morphological and metabolic changes of gliosis^41,43,68–70^. Both cell populations progressively enveloped the lesion, forming the outside layers of the glial scar (Fig. 3, Supp. Fig. 5), and expressed common reactivity markers (*Gfap, Serpina3n, C4b*; Fig. 4-5, Supp. Fig. 4). The core reactivity signature is shared in glial populations across pathologies including AD^36,37,44–46^, multiple sclerosis^41,42^, aging^43^, spinal cord injury^24,30,67,71^ and recently ischemic stroke^40,72^, indicating conserved aspects of activation^34,69^. In accordance, our reference-free deconvolution identified injury-related topics of expression programs enriched for profiles of disease-associated astrocytes and of oligodendrocytes (Fig. 3). Notably, the topics were defined by markers of immature and injured astrocytes (*Igfbp2* and *Fabp7*)^30,36,39^, genes involved in pathology response of oligodendrocytes (*Klk6, Trf*)^30,54,59^, but also microglia (*Hexb, Csf1r, Trem2, Cst7*)^37,40,45^, further highlighting injury-induced response of multiple cell type components. An interesting pattern, clearly observable on day 7, is the extension on the reactive topics to white matter tracts and striatum (Fig. 3-5, Supp. Fig. 7 and 8). This suggests spreading of the pathophysiological changes out of the cortex, potentially representing early signs of secondary neurodegeneration frequently observed in ischemic patients^73^.

The role of oligodendrocytes in acute CNS injuries and neurodegeneration remains elusive. We recovered three trauma-associated populations of oligodendrocytes that appear to contribute to the management of the injury. On day 1, MOL DA2 was the most abundant injury-induced population, accumulated near the lateral ventricle (Fig. 5). MOL DA2 had increased energy demand and enhanced translational activity including expression of genes influencing survival, factors indicating a role of ancestor to other injury-induced populations or being involved in their priming. With the highest levels of *Serpina3n* and *Klk6*, MOL DA1 is likely the main source of anti-inflammatory response. Overexpression of *Serpina3n* protease inhibitor was shown to reduce apoptosis and neuroinflammation, and ultimately stimulate cell proliferation through the activation of the Akt-mTOR pathway^60^. The population of MOL DA1 also overexpressed *Klk6*, which is an extracellular protease acting against myelin production^74^. Premature myelination has recently been shown to inhibit regrowth of neurons^75^, pointing to a potential regulatory mechanism supporting neuronal reorganization after the injury. Post injury, the MOL IFN population gradually strengthened its signature near the lesion core. Similar interferon-responsive populations were recently identified for microglia^43,45^ and astrocytes^76,77^, suggesting these populations crosstalk actively regulating the inflammatory status of the lesion.

There are several limitations to consider in this work. For practical reasons, our study has limited design, being performed only on young male mice screened for the first week of injury. Although both the age- and sex-related risk factors are important subjects for further understanding of stroke mechanisms and outcome^2^, they need to be analyzed in dedicated studies. Similarly, analysis of later time points holds great promise of providing valuable insight into processes related to chronic neuroinflammation, its role in patient recovery and for the onset of other neurodegenerative diseases. Despite these limitations, this work provides a complex reference atlas of the experimental ischemic brain injury, allowing for further data mining and hypothesis testing.

We have presented a comprehensive transcriptomic study providing insights into molecular and cellular processes in the acute phase of experimental ischemic brain injury, with temporal, spatial and single-cell resolution. This work represents an important resource for further studies aiming to elucidate stroke pathology and may contribute to the search for new targets for treatment.

## Methods

### Mice

All procedures involving the use of laboratory animals were performed in accordance with the European Communities Council Directive 24 November 1986 (86/609/EEC) and animal care guidelines approved by the Institute of Experimental Medicine, Academy of Sciences of the Czech Republic (Animal Care Committee on March 15, 2022; approval number 50/2020). All efforts were made to minimize both the suffering and the number of animals used. Experiments were performed on 3-month-old C57Black/6 male mice. In addition, for single-cell RNA-seq and immunohistochemistry we used B6.Cg-Tg(Plp1-cre/ERT)3Pop/J which was cross-bred with B6;129S6-Gt(ROSA)26Sortm14 (CAG-tdTomato)Hze/J (Jackson Laboratory, Bar Harbor) also termed Plp1/tdTomato mouse. The expression of tamoxifen-inducible Cre recombinase is controlled by the Plp1 promoter and after tamoxifen administration, tdTomato red fluorescent protein is expressed in Plp1 positive cells, which includes oligodendrocytes. Tamoxifen was administered intraperitoneally for two days (100 mg/kg, Sigma–Aldrich)^78^. The mice were kept on 12-hr light/dark cycles with access to food and water *ad libitum* and assigned randomly to experimental groups.

### Middle cerebral artery occlusion (MCAO), a model of experimental stroke

Prior to the induction of MCAO, mice were anaesthetized with 3% isoflurane (Aerrane, Baxter) and maintained in 2% isoflurane using a vaporizer (Tec-3, Cyprane Ltd.). A skin incision between the orbit and the external auditory meatus was made, and a 1-2 mm hole was drilled through the frontal bone 1 mm rostral to the fusion of the zygoma and the squamosal bone, about 3.5 mm ventrally to the dorsal surface of the brain. The middle cerebral artery (MCA) was exposed after the dura was opened and removed. The MCA was occluded by short coagulation with bipolar tweezers (SMT) at a proximal location, followed by transection of the vessel to ensure permanent occlusion. During the surgery, body temperature was maintained at 37 ± 1°C using a heating pad. This MCAO model yields small infarct lesions in the parietal cortical region. Intact cortical tissue from 3 months old mice was used as control. We have evaluated the impact of sham surgery (same surgical procedure, without coagulation of MCA) on gene expression in a previous study Androvic et al.^72^, which we did not find differences between naive and sham-operated mice.

### Brain dissection

Mice were deeply anesthetized with pentobarbital (PTB) (100 mg/kg, i.p.), and perfused transcardially with cold (4–8°C) isolation buffer containing (in mM): NaCl 136.0, KCl 5.4, HEPES 10.0, glucose 5.5, osmolality 290 ± 3 mOsmol/kg. The forebrain hemisphere was isolated by dissection of the olfactory lobes and the midbrain structure (bregma from ∼ +3 to -4 mm).

### Magnetic Resonance imaging (MRI)

All *in vivo* imaging experiments were collected on a 4.7 T MR scanner (Bruker BioSpec, BioSpin, Ettlingen, Germany) equipped with an optimized ^1^H radiofrequency surface coil custom‐designed and constructed in a magnetic resonance laboratory. Five mice with MCAO-induced lesions were scanned per timepoint (1, 3 and 7 days post injury, respectively), and one additional mouse underwent a repeated scanning at all of the three timepoints. During the scanning, the animals were anesthetized with 1.5% isoflurane mixed with air. The measurement protocol consisted of a standard 2D Rapid Acquisition with a Relaxation Enhancement (RARE) multi-spin echo sequence acquiring T_2_-weighted sagittal, coronal and axial images. For volumetry, coronal and axial planes were used with these basic imaging sequence parameters: repetition time (TR) = 3300 ms, effective echo time (TE) = 36 ms, excitation and refocusing pulses = hermite pulses, number of acquisitions (NA) = 8, scan time (ST) = 11 min 52 s, turbo factor = 8, slice thickness = 0.6 mm, field of view (FOV) = 35 mm and spatial resolution = 137 × 137 µm^2^. The volumetry of lesioned area and its 3D reconstruction were assessed manually using VGstudio Max (v2.1, Volume Graphics) in each axial and coronal T_2_-weighted image (the average from these two planes was calculated). Wilcoxon rank sum test was used for statistical testing of the lesion volume difference. For 3D reconstruction, two stacks of images covering whole mouse brain were acquired with slice thickness = 0.4 mm, ST = 17 m 48s, NA = 12. All other imaging parameters were identical.

### 10X Visium Spatial transcriptomics

#### Tissue preparation and sectioning

Dissected forebrains were embedded in optimal cutting temperature (OCT) medium, rapidly placed on dry ice to freeze, and then transferred to -80°C for a maximum of six weeks of storage. The forebrains were sectioned coronally (10 µm thickness) using Leica CM1950 cryostat (Leica Microsystems). Sections collected for 10X Visium Spatial Gene Expression processing were stored at -80°C.

#### Section fixation, staining and imaging

Fixation, staining, and imaging was performed strictly according to manufacturer’s manuals as in “Methanol fixation + H&E Staining Demonstrated protocol” (CG000160) and “Imaging Guidelines Technical Note” (CG000241). In brief, the sections were shortly incubated to thaw, methanol-fixed, isopropanol-incubated, and H&E stained. Stained sections were imaged using Carl Zeiss AxioZoomV16 upright microscope equipped with PlanNeoFluar Z objective (2,3x magnification, 0,57 NA, 10,6 WD) at total 63,0x zoom. Images were imaged with 5% overlap and stitched using ZEN blue pro 2012 software.

#### Sequencing library preparation

Sequencing libraries were prepared according to the manufacturer’s manual “Visium Spatial Gene Expression Reagent Kits User Guide” (CG000239). In brief, directly after imaging, the sections were permeabilized for 9 minutes, which is the optimal time determined in the “Visium Spatial Gene Expression Reagent Kits - Tissue Optimization User Guide” (CG000238) experiment. When permeabilized, on-slide reverse transcription was performed followed by second strand cDNA synthesis. Next, the double-stranded cDNA was transferred to a microtube, PCR-amplified, enzymatically fragmented, size selected, and tagged with Illumina sequencing adapters. The quality of each library was assessed by capillary electrophoresis on FragmentAnalyzer using the NGS High Sensitivity kit (Agilent, DNF-474). Sample libraries were pooled and sequenced on Illumina NovaSeq 2000 targeting 50,000 read pairs per tissue-covered spot.

#### Low-level data processing

The raw sequencing data were processed using the recommended set of *Spaceranger* function (v1.2.2, 10X Genomics) for processing of fresh frozen samples. Binary base call files were demultiplexed using *mkfastq* function with default parameters. The resulting fastq files were mapped separately to spaceranger reference (mm10/GRCm38) using *count* function, which takes a microscope slide image and fastq files, performs alignment, tissue detection, fiducial detection, and barcode/UMI counting.

#### Data processing and clustering – control section

The analysis was performed using R-based *Seurat* pipeline (v4.1.0, https://github.com/satijalab/seurat)^12^, accessible on LabGenExp Github). First, we counted quality metrics, excluded spots with less than 100 unique genes from the analysis and normalized the counts with the *SCTransform* function using the maximal number of identified variable features. We performed principal component analysis (PCA) for the top 100 principal components (PCs) on the genes of nuclear origin. Determined the optimal number of PCs by inspecting their standard deviations (*ElbowPlot* analysis) and spot PC scores (*DimHeatmap* analysis) to determine the variance they capture. We used the top 25 PCs as input for the UMAP dimensionality reduction. Next, we determined the clusters using functions *FindNeighbors* and *FindClusters* with the Leiden algorithm for modularity optimization. We examined the clusters at varying resolutions (from 0.6 to 2.0), visually identifying the robustness in the UMAP and Spatial plots. We used a resolution of 1.2, yielding 19 clusters, as baseline, which produced similar annotations to the Allen Brain Atlas (https://mouse.brain-map.org/static/atlas, images 64 and 65, Reference Atlas version 2, 2011)^13^. Additionally, the anatomically related clusters were merged, yielding a final 13 distinct brain region-specific clusters.

#### Region annotation and comparison to Allen Brain Atlas

We calculated differentially expressed genes (DEGs) for the individual brain regions (i.e., region vs. rest of the section) using the *FindAllMarkers* function. The DEG was considered as a brain region marker, when passing the criteria of log_2_-fold change > 0.58 and adjusted p-value < 0.01. Top 100 regional Allen Brain Atlas (ABA) *in-situ hybridization* markers (fold-change sorted) were obtained for each region separately (https://mouse.brain-map.org/search/index). The significances of overlaps between our top 100 log_2_-fold change sorted and top 100 ABA markers were calculated using hypergeometric test (*phyper* function, *stats* R package, v 4.2.2) with the full gene list of corrected count matrix as background set (n = 18302 genes). P-values were corrected for multiple hypotheses testing by the Benjamini-Hochberg procedure (false-discover rate, FDR).

#### Data processing and clustering – all sections

Spatial data were collected in two batches: Batch 1 consisted of a control (bregma -1.3 mm) and post-ischemic day 7 sections (bregma +0.5 mm); Batch 2 were three post-ischemic sections collected on day 1, day 3 and day 7 (bregma -1.3 mm). We filtered the low-quality spots (> 200 unique genes), normalized the datasets separately by batches (*SCTransform*), identified the maximal number of integration features (*SelectIntegrationFeatures*), computed the missing sctransform residuals (*PrepSCTIntegration*), identified the integration anchors (*FindIntegrationAnchors*), and merged the two datasets (*IntegrateData*). Using the matrix of centered, corrected Pearson residuals we calculated the top 100 PCs using the list of genes of nuclear origin. Using *ElbowPlot* and *DimHeatmap* analyses, we chose the top 30 PCs as being most informative and used them as input for the UMAP dimensionality reduction. Clusters were determined using the *FindNeighbors* and *FindClusters* functions. We examined clustering robustness at varying resolution parameter values (0.6 to 2.0) and settled for the resolution of 1 yielding 27 clusters, which we annotated into 16 anatomically related brain regions. Next, we repeated the integration process as outlined above for the spots originating from the regions of the cortex and from the ischemic lesion. After data merger, top 20 PCs were used in the UMAP clustering, yielding 19 clusters. Out of these six defined injury-induced clusters. Finally, we combined clustering from the whole-section and cortical integrations, resulting in the final annotation of 16 brain regions and 6 ischemic areas. For the final presentation, we excluded the post-ischemic day 7 section (bregma +0.5 mm), due to its different localization. The section was, however, included in the integration, to allow for mutual anchoring of lesion clusters. Data from the removed section are part of the publicly available deposited dataset.

#### Pathway enrichment

Using the *FindAllMarkers* function we calculated DEGs for each annotated region (log_2_ fold-change threshold > 0.58 and adjusted p-value < 0.01). On the list of marker genes, we performed the Gene Ontology (GO) analysis of biological processes, cellular components, and molecular functions, using the *enrichGO* function (*clusterProfiler* package v4.2.2), with the 18302 genes as background and FDR-adjustment of p-values (FDR < 0.01). For pathway enrichment we used Metascape^27^ individually for each section to generate the networks of commonly enriched terms in the ischemic regions. We recapitulated the enrichment pathway settings, searching for GO biological processes with minimal overlap of five genes at a significance level of <0.01. The network maps were generated automatically with the Metascape tool, for which we reviewed the cluster annotation based on the GO terms included within the parent cluster.

#### Re-analysis of the Milich et al. single-cell dataset

We obtained the Milich et al.^30^ dataset from GEO omnibus (GSE162610) and re-analyzed it using R-based *Seurat* pipeline (v4.1.0, https://github.com/satijalab/seurat)^12^. Data consisted of samples collected in three batches. Upon data quality control, we discarded the first and third batch, which did not contain all the sampled cell types and/or had low transcript coverage. Using batch 2 data only, we filtered the set for cells with >200 unique transcripts, and normalized the data (*SCTransform* using the maximal number of identified variable features). We computed PCA for the top 100 PCs on the list of genes of nuclear origin. Using the *ElbowPlot* and *DimHeatmap* analyses, we identified the top 30 PCs accounting for suffient variation in the data and used them for UMAP reduction. Clusters were calculated using *FindNeighbors* and *FindClusters* functions with the Leiden algorithm for modularity optimization at a resolution of 0.8. The procedure yielded 31 clusters, which we annotated to 17 different cell types, considering their marker expression (*FindAllMarkers*, log_2_FC > 0.58, padj < 0.01) and the original annotation^30^.

#### Performing RCTD deconvolution

We performed the Robust Cell Type decomposition (RCTD) deconvolution using the *spacexr* R package v2.0.0. We prepared the *Reference object* from the raw count matrix of the in-house re-analyzed batch 2 of Milich et al.^30^ dataset. For the spatial data, we prepared a separate *SpatialRNA object* from each section separately. *SpatialRNA objects* were similarly prepared from the raw RNA count matrix (‘*Spatial’* assay in the *SeuratObject*) and their spot coordinates. Next, we created a separate *RCTD object* for every section using *create*.*RCTD* with the *Reference object* and *SpatialRNA object* as inputs. We deconvoluted the *RCTD objects* using *run*.*RCTD* with the doublet mode parameter set to ‘multi’ to allow fitting the number of cell types per spot. This resulted in a matrix of ratios of cell type levels per spot.

#### Reference-free deconvolution – STdeconvolve

For reference-free restoration of cell type signatures from spatial data we used *STdeconvolve*, an R package developed by Miller et al.^51^. *STdeconvolve* is a reference-free, unsupervised algorithm built on latent Dirichlet allocation, originally suited for discovery of latent topics (here cell type signatures) in collections of documents. We ran *STdeconvolve* v0.99.12 on each section separately, first creating a gene count matrix and spot coordinates filtered for genes with >10 reads and spots with >100 detected genes. We identified genes with more-than-expected variability using the *restrictCorpus* function, defined by the presence in-between 5 to 95 % of spots at over-dispersion parameter alpha at the default 0.05 value. Next, the LDA models were fitted using *fitLDA* function with default settings, screening models in a range of 15 to 30 topics (the K parameter). The optimal number of topics was evaluated with respect to the number of rare cell type signatures (overall presence <5 %) and the perplexity score. Using this approach, we identified 23, 24, 24 and 21 topics in control, day 1, day 3 and day 7 sections, respectively. Result was a matrix of topic ratios for each spot (≈ cell type signature abundancies), as well as quantitative measures of cell-type gene probabilities on per topic basis (≈ topic characterization via estimated gene expression).

#### Signature expression

To visualize a module of common genes, either as a set of population markers or a biological process, we used *AddModuleScore* function (*Seurat* package), which compares the average expression of the signature genes with permuted randomly selected genes of similar expression levels (size = 100 genes, 24 permutations). To visualize populations described in this study, we used top 15 markers (>0.58 log2FC sorted, P_adj_ < 0.01) for the populations described in this study, and full marker gene lists for the published populations. To visualize GO terms, we used all its DEGs that passed the marker criteria (log_2_-fold change > 0.58 and adjusted p-value < 0.01).

### Bulk RNA-Seq

#### RNA isolation, library preparation and sequencing

Samples were collected from the injured cortical region (including affected as well as non-affected brain tissue) and homogenized using the TissueLyser (QIAGEN). Total RNA was extracted with TRI Reagent (Sigma-Aldrich) according to the manufacturer’s protocol and treated with TURBO DNA-free kit (Thermo Fisher). RNA quantity and purity was assessed using the NanoDrop 2000 spectrophotometer (Thermo Fisher) and RNA integrity was assessed using the Fragment Analyzer (Agilent). All samples had RQN > 8. Libraries were prepared from 400 ng total RNA with QuantSeq 30 Library Prep Kit FWD (Lexogen) according to manufacturer’s protocol. 1 µl of ERCC spike-in (c = 0.01×; Thermo Fisher) per library was included. Libraries were quantified on the Qubit 2 fluorometer (Thermo Fisher) and Fragment Analyzer (Agilent), and sequenced on the NextSeq 500 high-output (Illumina) with 85 bp single-end reads. 11.5 – 38 million reads were obtained per library with a median of 16 million reads.

#### Data processing, mapping, and counting

Adaptor sequences and low-quality reads were removed using TrimmomaticSE v0.36^79^. Reads mapping to mtDNA and rRNA were filtered out using SortMeRNA v2.1 with default parameters^80^. The remaining reads were aligned to GRCm38 and ERCC reference using STAR v2.5.2b with default parameters^81^. Mapped reads were counted over Gencode vM8 gene annotation using htseq-count with union mode for handling of overlapping reads^82^.

#### Differential expression and pathway enrichment

We performed differential expression testing using DESeq2 v1.34.0^83^. We filtered the data for genes with >10 reads and normalized them by *varianceStabilizingTransformation* function. We applied the DESeq model using *DESeq* function with type of fit parameter set to ‘local’. Statistical testing was done by comparing the control sample with each timepoint individually. Genes with log_2_ fold-change >0.58 and adjusted p-value <0.01 were considered as differentially expressed markers. Overlap of the upregulated markers between the spatial dataset (made as union of the core and periphery markers) and bulk markers was tested for significance using hypergeometric test (*phyper* function, *stats* R package v 4.2.2). The gene set enrichment analysis of biological processes was performed on the *DESeq* output file using *clusterProfiler* R package (v4.2.2). Threshold for the significantly enriched biological processes was p-value <0.05 after the FDR adjustment.

### Single-nucleus RNA-Seq

#### Sample and sequencing library preparation

Injured forebrain hemispheres were frozen on dry ice and stored at -80 °C until processing. For each condition, three hemispheres were collected and pooled during the single-nucleus suspension preparation. All steps for nuclei isolation were performed on ice, with instruments and plastics pre-chilled on ice, or centrifugation steps at 4°C. The frozen brain tissues were combined with 2 ml of lysis buffer (10 mM Tris, pH = 7.4; 10 mM NaCl; 3 mM MgCl2; 0.1 % NP-40; 0.2 U/µl RNaseOUT; 0.32 M Sucrose; 1x protease inhibitor, Roche) and transferred to 2 ml Dounce tissue grinder (Sigma). Tissues were homogenized by 20 strokes with pestle A, followed by 20 strokes with pestle B and filtered using 30 µm filter (Celltricks). Tissue grinder was rinsed with 1 ml HEB buffer (Hibernate A, ThermoFisher; 10 µl/ml GlutaMAX, ThermoFisher; 0.32 M Sucrose) and the wash filtered into the same tube. Homogenates were split by careful pipetting of 1.2 ml into two 2 ml tubes prefilled with 500 µl sucrose cushion (1.2 M sucrose; 10 mM Tris; 10 mM NaCl; 3 mM MgCl2; 0.2 U/µl RNase OUT) and centrifuged at 13000 g for 20 minutes at 4°C. Supernatants were removed, except 100 µl to keep the pellets covered. Pellets with the remaining supernatant were resuspended in 1 ml sucrose solution (1.0 M sucrose; 10 mM Tris; 10 mM NaCl; 3 mM MgCl2; 0.2 U/µl RNase OUT), pipetted into 2 ml tubes prefilled with 500 µl sucrose cushion and again centrifuged at 13000 g for 20 minutes at 4°C. Supernatants were removed and the pellets resuspended in 1 ml of Nuclei Wash Buffer (DPBS without calcium and magnesium; 0.5 % BSA; 0.2 U/µl RNase OUT; all ThermoFisher) and transferred into 1.5 ml tubes. Samples were centrifuged at 500 g for 5 minutes, the pellets resuspended in 1 ml of Nuclei Wash Buffer and centrifuged again. The final pellets with nuclei were resuspended in 1 ml of Nuclei Wash Buffer and filtered using 30 µm filter. Nuclei stained with Propidium Iodide were counted using CellDrop Automated Cell Counter (Denovix) at default settings. Nuclei suspensions were diluted to final concentrations of 1000 nuclei/µl and single-nucleus RNA-Seq (snRNA-Seq) libraries were prepared using Chromium Next GEM Single Cell 3’ Kit v3.1 (1000268, 10X Genomics) according to the manufacturer’s protocol. Final libraries were eluted with EB buffer with 0.1% Tween, and their quality was assessed by capillary electrophoresis on the FragmentAnalyzer using the NGS High Sensitivity kit (DNF-474, Agilent). The libraries were pooled and sequenced on an Illumina NovaSeq 2000. Approximately 100M reads per sample were obtained.

#### Data processing

Libraries were controlled for contamination using fastq_screen v0.11.1. Reads were mapped against the mouse genome GRCm38 and counted using annotation genecode.vM8 using STAR v2.7.3a with parameters “--soloType CB_UMI_Simple -- soloFeatures GeneFull –soloCBmatchWLtype 1MM_multi_pseudocounts –soloUMIdedup 1MM_Directional –soloUMIfiltering MultiGeneUMI”. Empty droplets were filtered out using *emptyDrops* command from *DropletUtils* package (v 1.14.1)^84^ with lower parameter set to 1000. All droplets with FDR < 0.001 were considered as nuclei. UMI count matrix was processed using the *Seurat* pipeline (*Seurat* v4.1.0, https://github.com/satijalab/seurat)^12^, accessible here: LabGenExp Github). Background RNA was removed using SoupX package (v 1.5.2)^85^ set at the default parameters. UMI counts were normalized using *SCTransform* and *Seurat* integrated. Nuclei were clustered using top 30 PCs and visualized using UMAP. In total, 26 clusters were identified and manually annotated based on the expression of known markers. Next, neuronal and non-neuronal cells were separated and processed for detailed annotation. Cells across the samples were merged, *SCTransform*-ed, clustered and re-annotated into the major populations.

#### Data analysis

Three populations were selected from non-neuronal populations, astrocytes (including ependymal cells and neuroblastema cells), microglia, and oligodendrocyte-lineage cells, and analyzed them separately using the *Seurat* pipeline (LabGenExp Github). UMI counts were normalized, *SCTransformed*, clustered and annotated. DEGs were calculated (*FindAllMarkers* function) and population markers defined (DEGs with log_2_ fold-change threshold > 0.58 and adjusted p-value < 0.01). On the list of marker genes, we performed the Gene Ontology (GO) analysis of biological processes, cellular components, and molecular functions, using the *enrichGO* function (*clusterProfiler* package v4.2.2), with the size of full gene list of the normalized matrix as for the background set, and selected the significant process based on the FDR-adjusted p-values (FDR < 0.01). Signature expression of the identified populations were visualized in the spatial sections as described in the *Spatial transcriptomics – Signature expression* Method’s section.

#### Metadata analysis

Markers of the published populations were obtained from the respective materials provided by the study’s authors. Signature expressions were calculated using *AddModuleScore* function (*Seurat* package), which compares the average expression of the signature genes with a permuted randomly selected genes of similar expression levels (size = 100 genes, 24 permutations). The entire signature gene list was used to calculate the signature. To calculate marker overlaps, the obtained list of markers was filtered for genes present in the normalized SCT matrix, and then significance tested using the hypergeometric test (*phyper* function), with the full gene list as background set.

### Single-cell transcriptomics

#### Sample preparation

The Plp1/tdTomato mice were deeply anesthetized with pentobarbital (PTB) (100 mg/kg, i.p.), and perfused transcardially with a cold (4–8°C) isolation buffer containing (in mM): NaCl 136.0, KCl 5.4, HEPES 10.0, glucose 5.5, osmolality 290±3 mOsmol/kg. To isolate the control and ischemic areas, the brain was sliced into 600 µm coronal sections using a vibrating microtome Leica VT1200S (Leica Microsystems). The collected tissue was incubated with continuous shaking at 37°C for 45 min in 1 ml of papain solution (20 U/ml) and 0.2 ml DNase (both from Worthington) prepared in isolation buffer. After papain treatment, the tissue was mechanically dissociated by gentle trituration using a 1 ml pipette. The dissociated cells were layered on top of 5 ml of ovomucoid inhibitor solution (Worthington) and harvested by centrifugation (70 x g for 6 min). This method routinely yielded ∼2 × 10^6^ cells per mouse brain. Cell aggregates were removed by filtering with 70 µm cell strainers (Becton Dickinson). Three animals per condition were pooled in the preparation of cell suspension. After obtaining an aliquot for FACS, the cell suspension was spin down, concentrated, and used for library preparation.

#### Collection of single cells

Single cell suspension from Plp1/tdTomato mice were sorted using fluorescent activated cell sorting (FACS; BD Influx). The flow cytometer was manually calibrated to deposit a single cell in the center of collection tube. Hoechst 33258 (Life Technologies) was added to the suspension of cells to check viability. 3000 Plp1+ cells from each group (control, D1, D3, D7) were collected into 96-well plates (Life Technologies) coated with BSA (ThermoFisher Scientific) and containing 5 μl of DMEM (ThermoFisher Scientific) and 15% FBS (HyClone).

#### Library preparation, data processing and analysis

Single-cell RNA-Seq (scRNA-Seq) libraries were prepared using Chromium Next GEM Single Cell 3’ Kit v3.1 (1000268, 10X Genomics) according to the manufacturer’s protocol. Final libraries were eluted with EB buffer with 0.1% Tween, and their quality was assessed by capillary electrophoresis on a FragmentAnalyzer (Agilent) using the NGS High Sensitivity kit (DNF-474). The libraries were pooled and sequenced on an Illumina NovaSeq 2000. Low-level data processing was performed similarly as for the snRNA-Seq data, with an exception in a STAR parameter “--soloFeatures Gene”. UMI count matrix was processed using *Seurat* pipeline (*Seurat* v4.1.0, https://github.com/satijalab/seurat)^12^. In brief, UMI counts were normalized (*SCTransform*), PCA-reduced, UMAP visualized, clustered (*FindNeighbors, FindClusters*), and annotated based on canonical markers. The dataset was then *Seurat* integrated with the single-nucleus counterpart. The analysis was further performed on the SCT-normalized corrected matrix, using identical set of tools as in the single-nucleus data Method’s section. For further details refer to LabGenExp github.

### Immunohistochemistry

#### Sample preparation and imaging

For immunohistochemical analyses, the animals were deeply anesthetized with PTB (100 mg/kg, i.p.) and perfused transcardially with 20 ml of saline followed by 20 ml of cooled 4% paraformaldehyde (PFA) in 0.1 M phosphate buffer. The brains were dissected out, post-fixed overnight with PFA, and treated with a sucrose gradient (ranging from 10% to 30%) for cryoprotection. Coronal 30-μm-thick slices were prepared using a cryostat (Leica CM1950, Leica Microsystems). For immunohistochemical staining, the slices were washed in a phosphate buffer saline followed by blocking of the non-specific binding sites with 5% Chemiblocker (Millipore), and 0.2% Triton in phosphate buffer saline. The blocking solution was also used as the diluent for the antisera. The slices were incubated with the primary antibodies overnight, and the secondary antibodies were applied for 2 hours at 4-8°C. The following primary antibodies were used: mouse anti-GFAP, 1:300; coupled to Alexa 488 (Ebioscience), goat anti Iba1, 1:500 (Abcam), goat anti Pdgfrα, 1:500 (RnDsystem), rabbit anti Col1a, 1:500 (Abcam) and goat anti Serpina3n, 1:100 (Biotechne). The secondary antibodies were goat anti-rabbit IgG or goat anti-mouse IgG conjugated with Alexa Fluor 488, 594 or 647 (ThermoFisher). All chemicals were purchased from Sigma-Aldrich unless otherwise stated. An Andor Dragonfly 503 spinning disk confocal microscope and 20x oil objective was used for the immunohistochemical analysis. Tile scans (15×15) of one hemisphere with stacks of consecutive confocal images were taken at intervals of 2 μm. Stitching of the tile scans were performed automatically by acquisition software Fusion (Oxford Instruments) with 10 % overlap. Maximum z-projection images were made using a Imaris (Oxford Instruments).

#### Cell counting (PLP1 and SERPINA3N)

To determine the number of cells, confocal images (15000 µm × 15000 µm × 30 µm) were taken covering whole hemisphere from the brain coronal slices. Those were prepared from them Plp1/tdTomato control mice and from the mice 1, 3 and 7 days after the ischemia (3 animals from each group, 3 brain slices) and stained for the SERPINA3N antibody. The percentage of double positive cells for PLP1/tdTomato and SERPINA3N were counted in the region of ischemia (core and glial scar) and the corresponding area in the control mice. Statistical testing was done using Wilcoxon rank-sum test.

## Author contribution

DZ and PA performed the spatial, single-cell and single nucleus experiments, and data analysis. DZ wrote the manuscript. DK performed all the mice work and immunohistochemistry experiments. DJ performed the MRI measurements. MA, MK, and LV coordinated the work and proofread the manuscript.

## Conflict of interest

Author MK was employed at TATAA Biocenter AB. The remaining authors declare the research was conducted in the absence of any commercial of financial relationships that could be construed as conflict of interest.

## Data availability

All data and code required to replicate the analyses, including spaceranger output files, H&E images and additional files will be available for download at Mendeley data. For interactive exploration, the spatial data will be available in Spatial Omics DataBase (SODB, https://gene.ai.tencent.com/SpatialOmics/)^86^. Sequence data for all samples presented in this study will be available upon publication.

## Acknowledgements

We would like to thank Sarka Benesova, Zuzana Matusova and Ravindra Naraine for helpful assistance, discussions and manuscript reading.

## Financial support

This study was supported by Czech Science Foundation (23-05327S, 23-06269S) and institutional support (RVO 86652036).

## Supplementary Figures

Supplementary figures are enclosed in a separate file.

