## Supplemental Figures and Tables for "Spatiotemporal transcriptomic map of ischemic brain injury": SpatiotemporalMap_SuppFigures.pdf

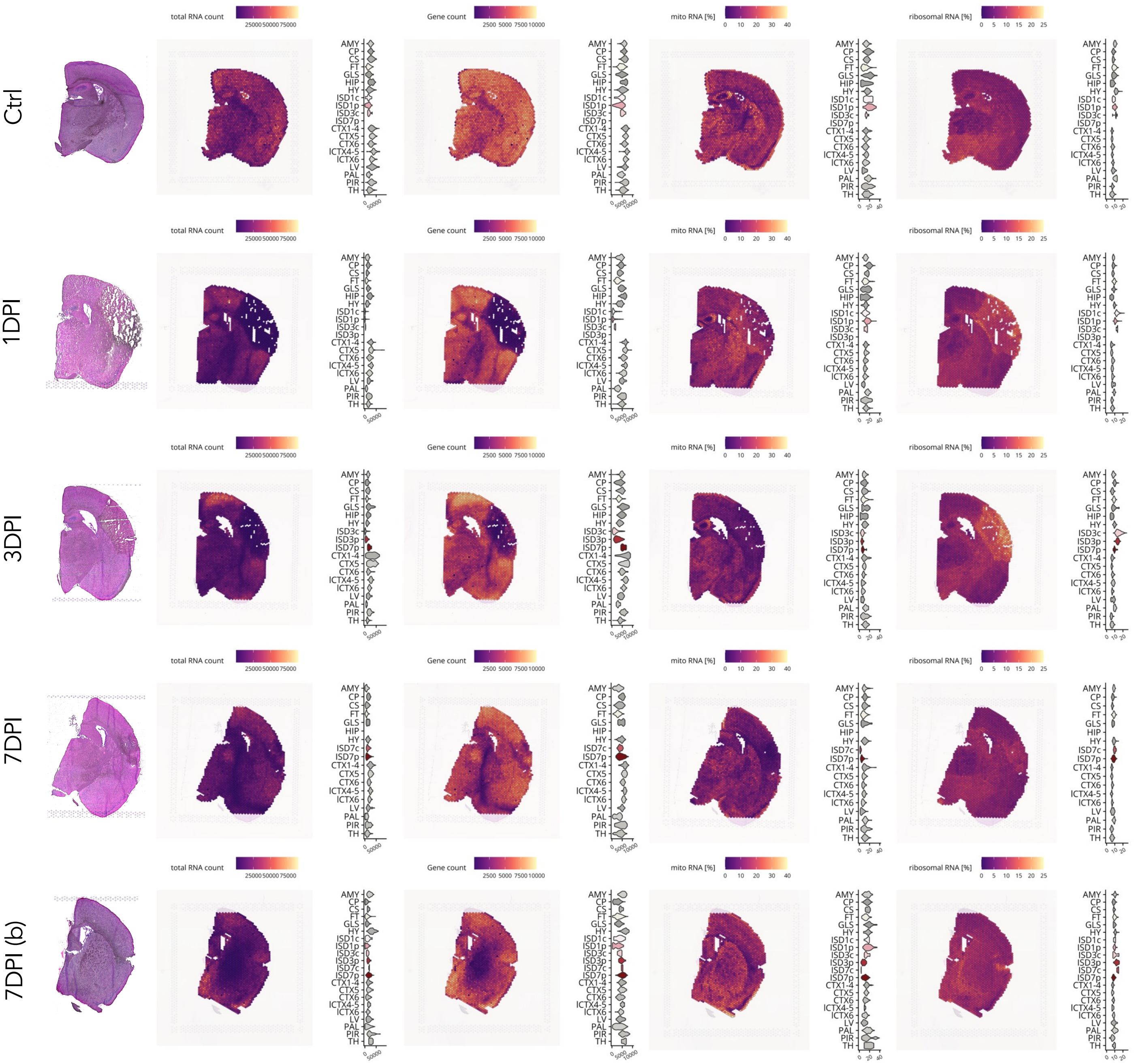


Supp. Figure 1 – **Quality control metrics of spatial transcriptomics dataset.** Hematoxylin-eosin (H&E) stained images of post-MCAO coronal mouse brain sections in the respective timepoints. The sections were collected from a location of bregma ~ -1.3 mm ± 0.1 mm, except the 7DPI(b) section of location bregma ~ +0.5 mm. The spatial plots show basic quality control metrics, including total unique molecular identifiers (UMI) count (total RNA count), gene count, and percentage of mitochondrial RNA (mito RNA, in %) and percentage of ribosomal RNA (in %). In the right-side violin plots, distributions of the metric values in the individual annotated brain regions are shown.

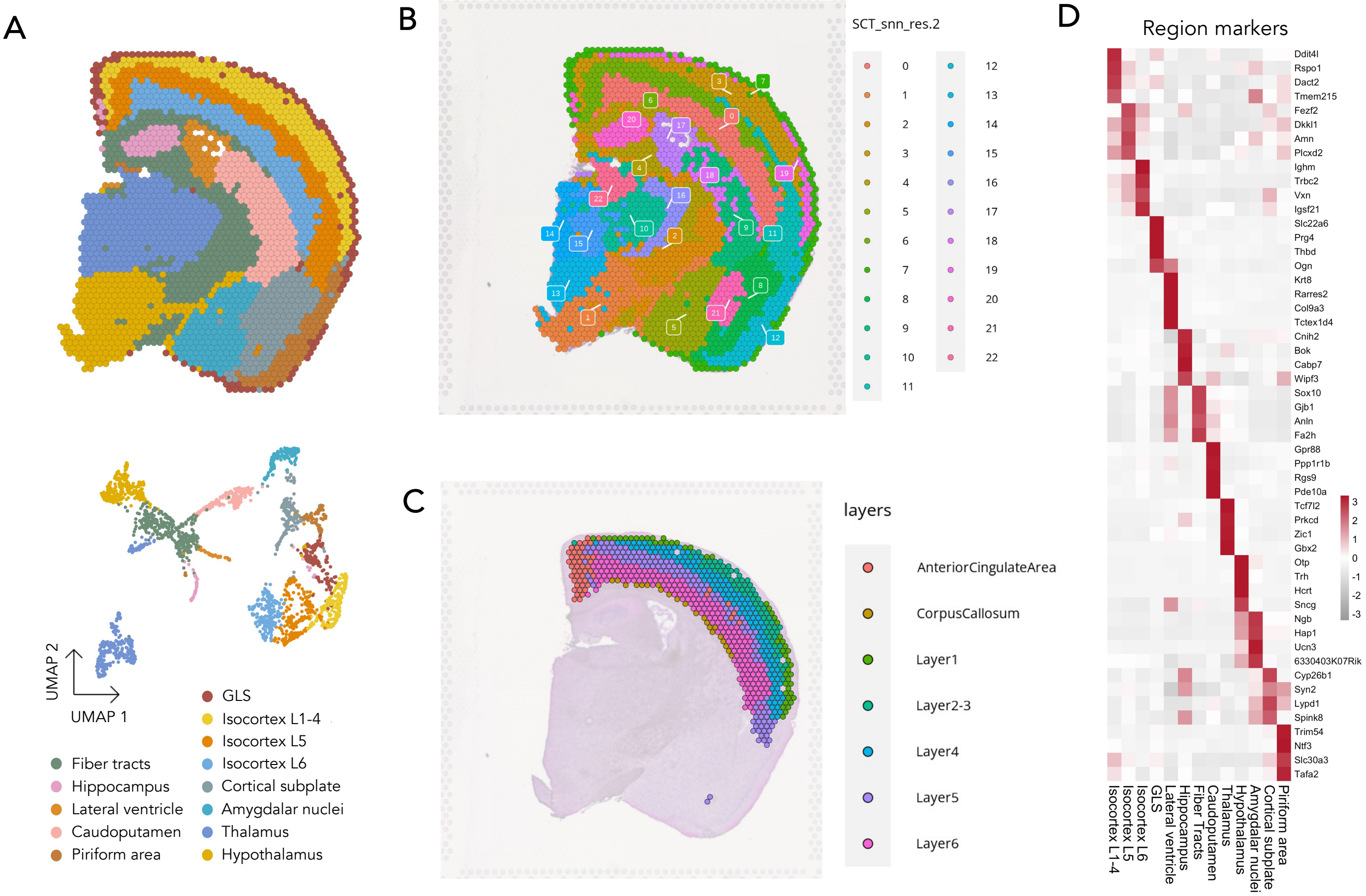


Supp. Figure 2 – **Spatial transcriptomics robustly capture underlying brain anatomy.** A) UMAP of spots, clustered according to their brain region of origin. B) Spatial plot of more detailed clustering, uncovering the more detailed brain regions, e.g thalamic reticular nucleus in cluster 16. C) Spatial plot of the cortical spots clustered according to the cortical layer of origin. D) Heatmap of markers identified for the brain regions of the control section. Normalized expression (color scale) is shown.

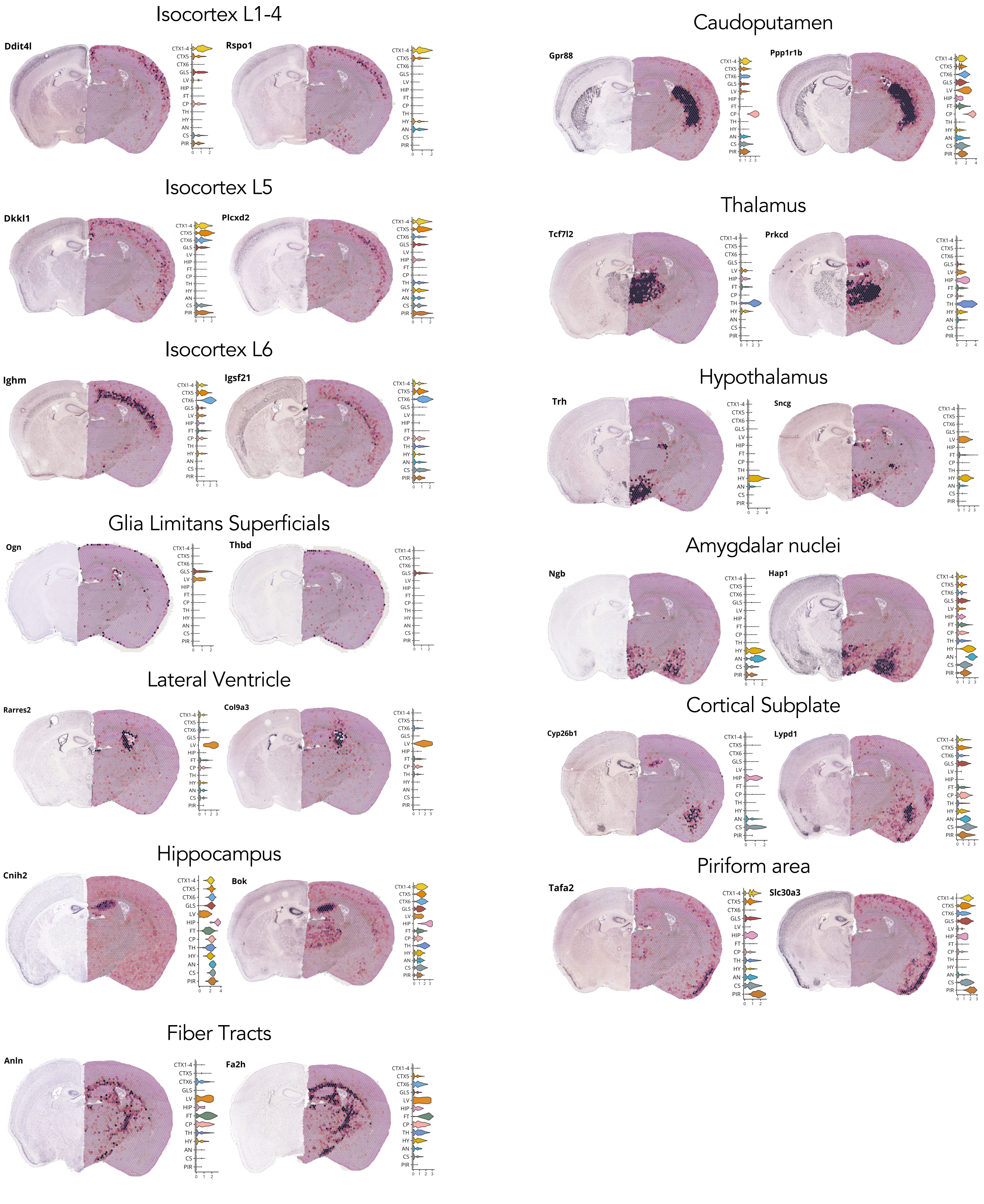


Supp. Figure 3 – **Region-specific expression of mouse brain markers**. Combined images of expression for selected markers, combining Allen Brain Atlas *in-situ* hybridization atlas (left, https://mouse.brain-map.org/static/atlas, images 64 and 65, Reference Atlas version 2, 2011, Sunkin et al 2013) and this study controls section (normalized expression). Distributions of the marker's normalized expression per brain region is shown in the right-side violin plots. The image locations correspond to bregma ~ -1.3 mm.

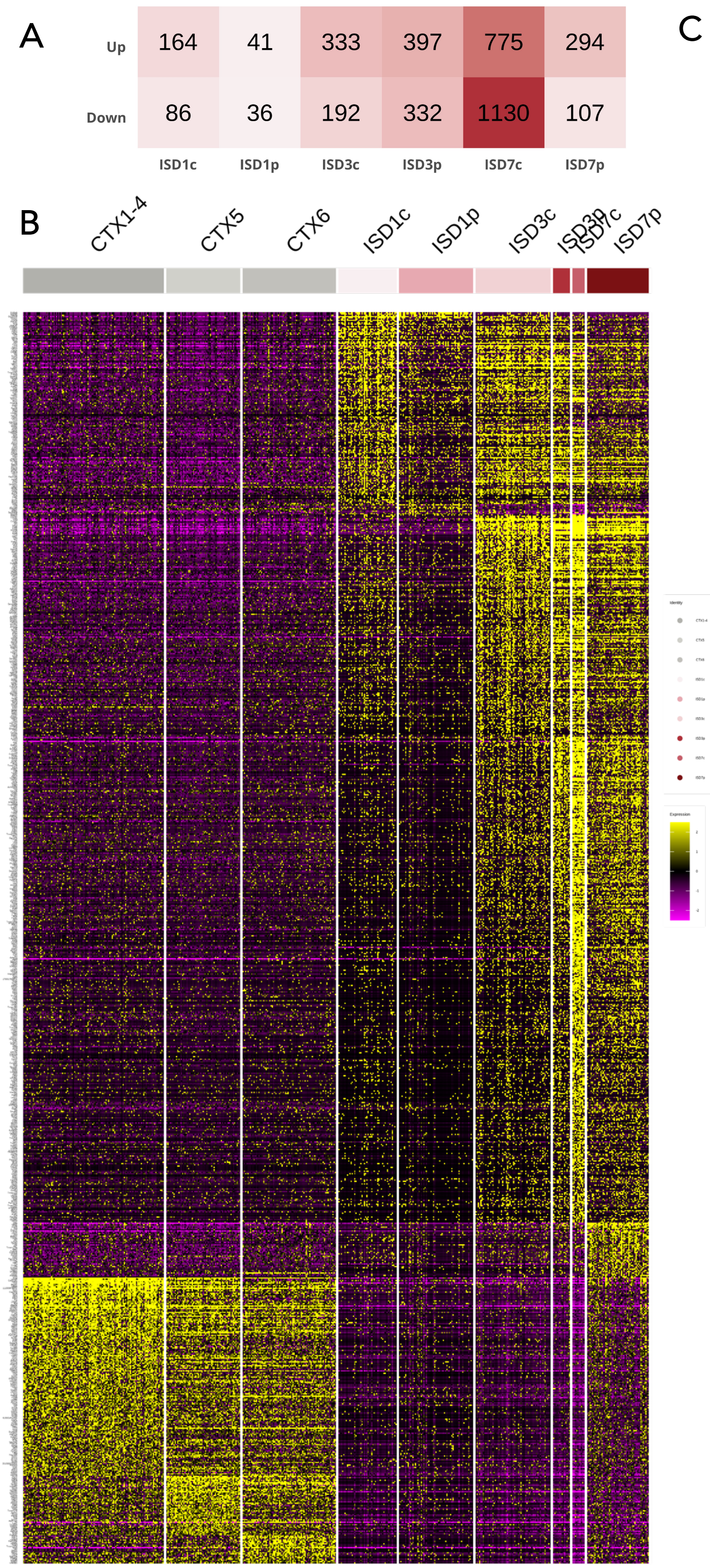


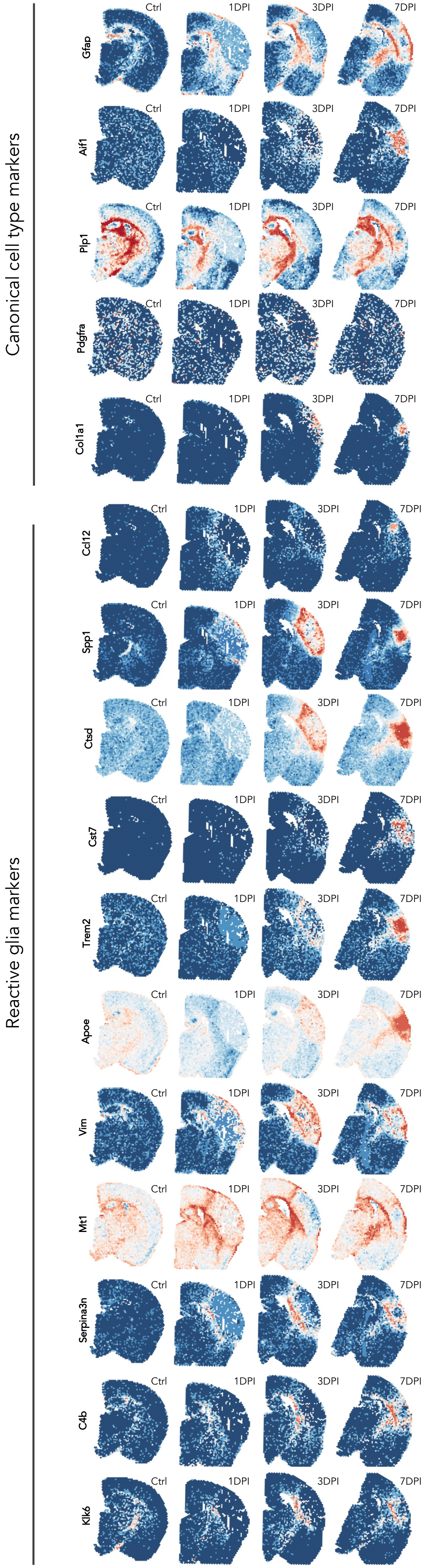


Supp. Figure 4 – **Markers of reactive glial populations localize to lesion periphery.** A) Differentially expressed genes (DEGs) in the lesion areas (Bonferroni-adjusted $P < 0.01$, $\log_2FC < -0.58$ for downregulated and $\log_2FC > 0.58$ for upregulated genes). B) Expression of the upregulated markers of the cortical regions visualized across 500 randomly sampled spots. C) Expression of selected markers of the glial populations and their reactive states.

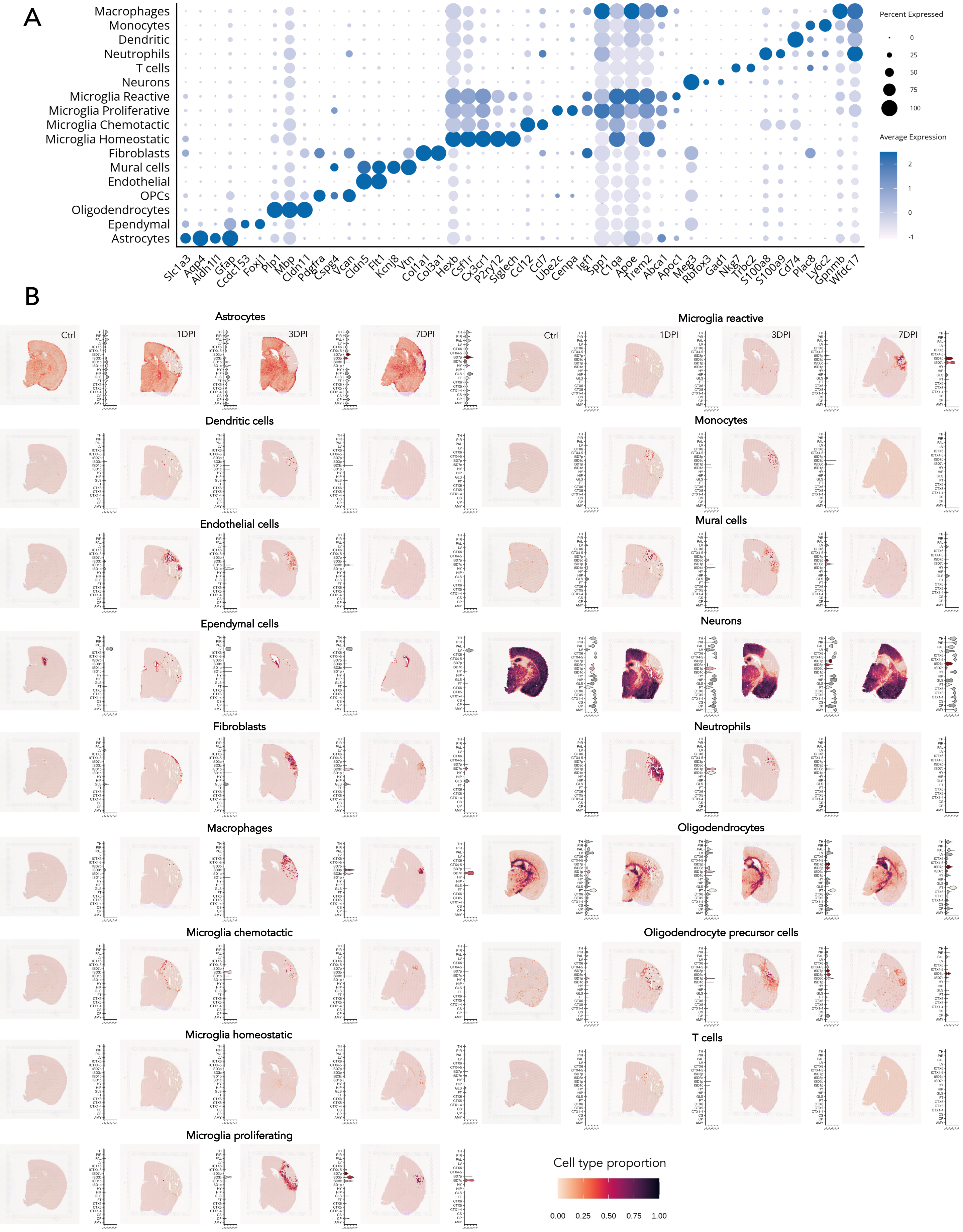


Supp. Figure 5 – **Cell populations identified in the re-analyzed Milich et al (2021) dataset used for RCTD deconvolution.** A) Expression of markers (color scale) for the cell populations identified after re-analysis of the Milich et al (2021) dataset, with the percentage of cluster's cells expressing the marker (dot size) are shown. B) Proportions of cell populations in the spatial sections (color scale), with the proportion distributions for the brain regions in the right-side violin plot are shown.

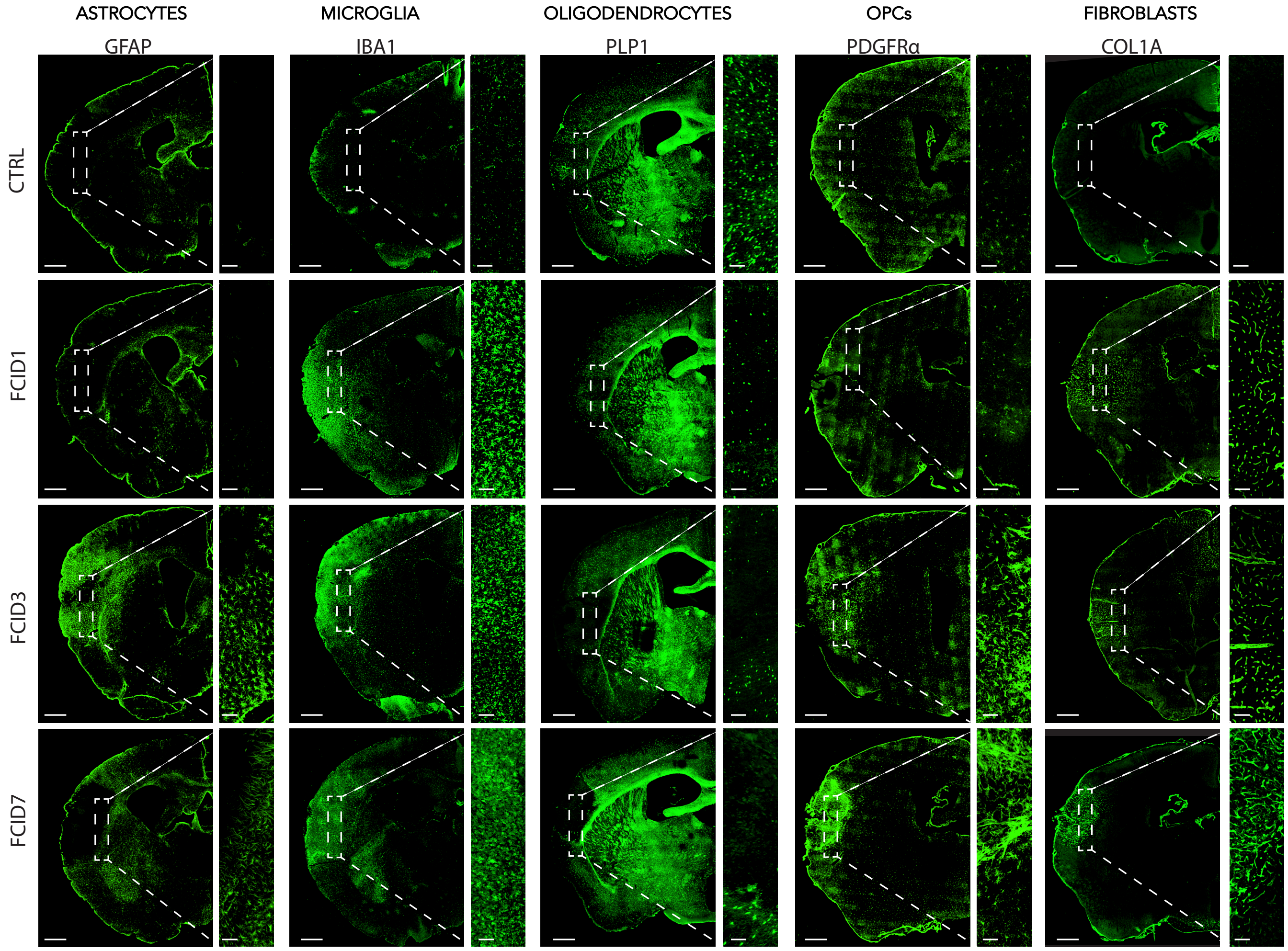


Supp. Figure 6 – **Validation of cell population presence using immunohistochemistry.** Coronal sections of the ipsilesional hemisphere following MCAO injury in mice, antibody stained with representative cell type markers. Scale bars: 500 μm for the hemisphere image, 100 μm for the inset image.

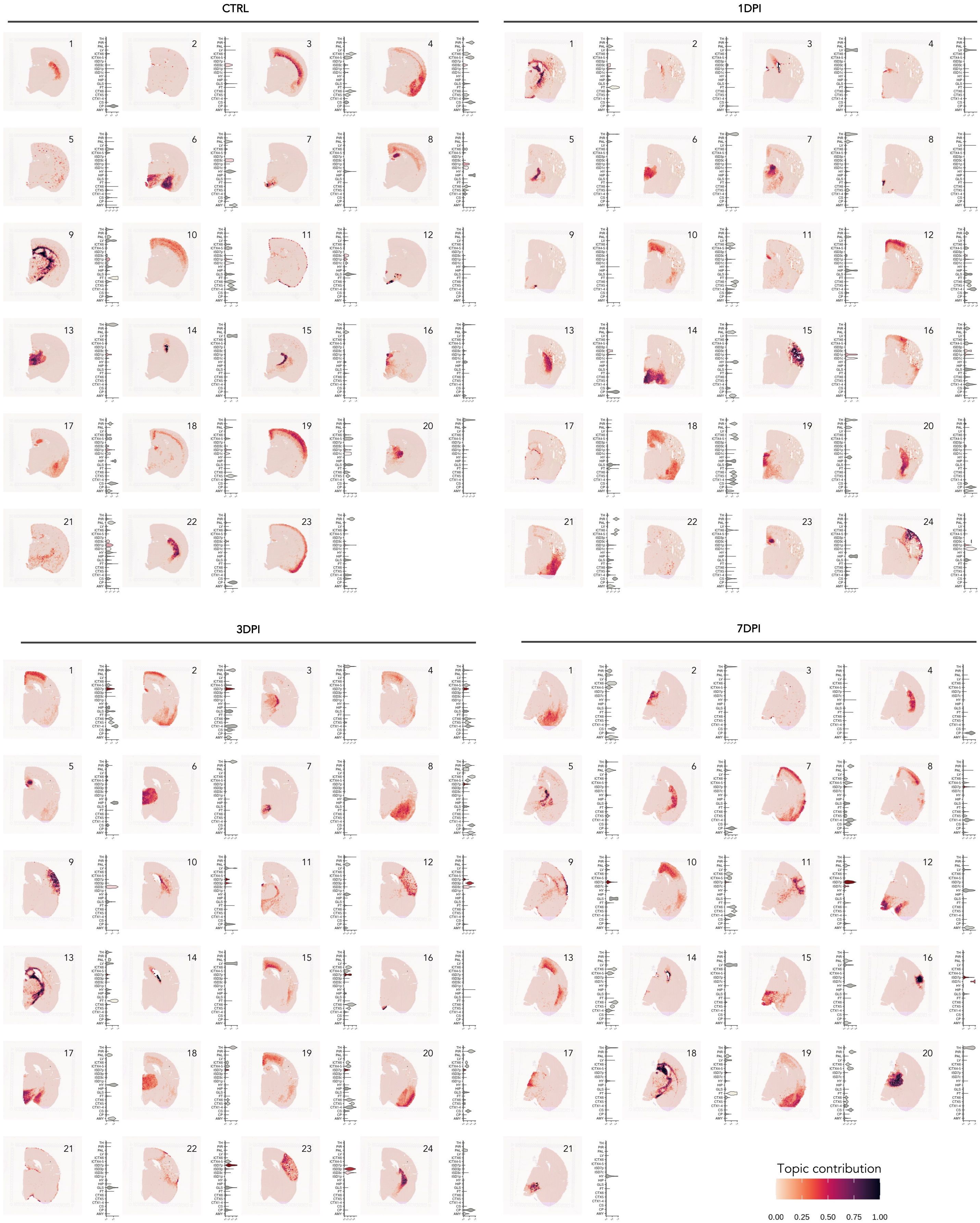


Supp. Figure 7 – **Identification of underlying expression modules (topics) using reference-free deconvolution algorithm (STdeconvolve, Miller et al 2022).** Topic contribution to the individual spots is shown (color scale). Per spot, topic contributions sum to 1. Distributions of the topic contributions in the individual brain regions are shown on the right-side. Topic marker genes are listed in Supp. Table 5.

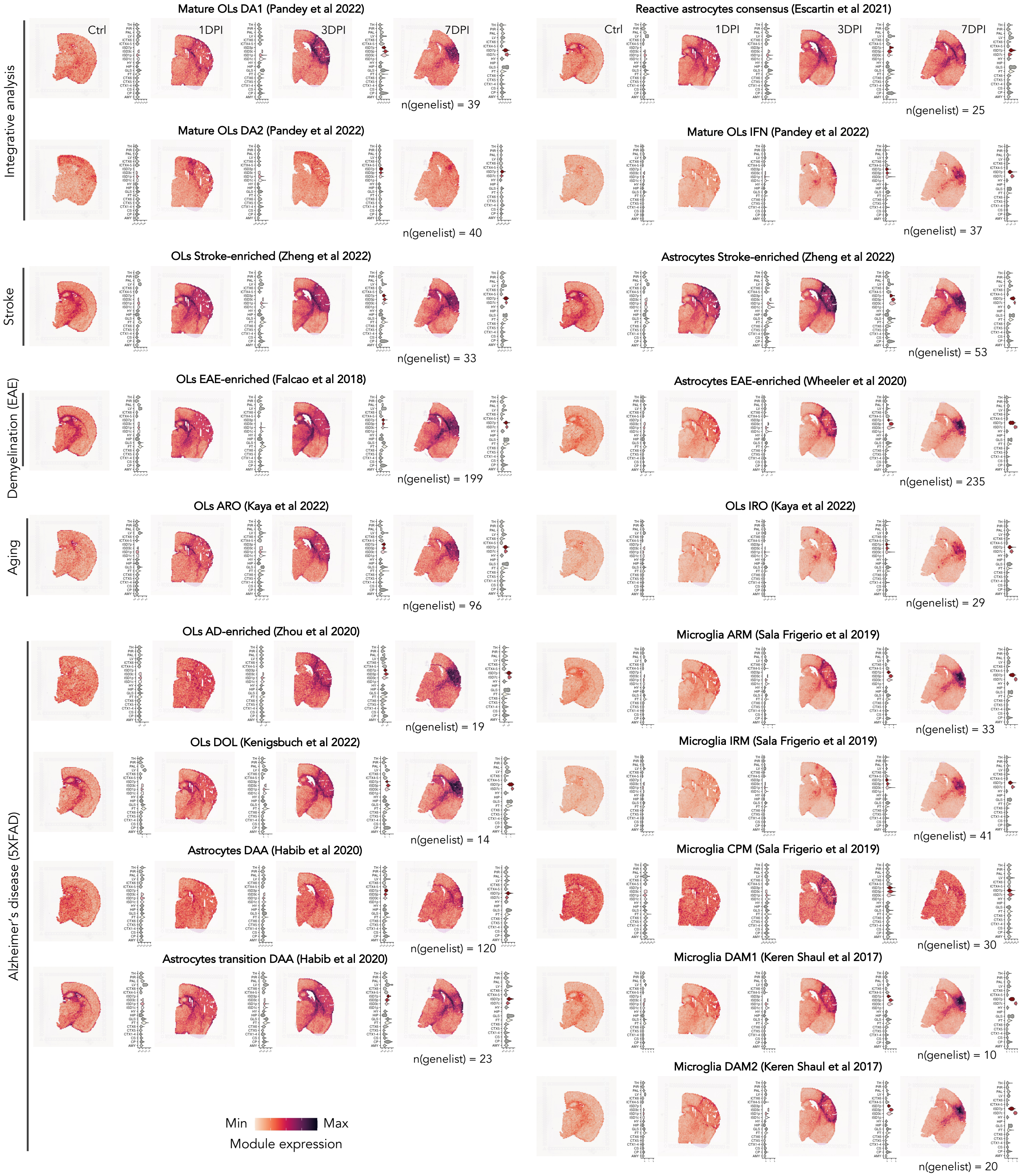


Supp. Figure 8 – **Metanalysis: Expression signatures of published reactive glia populations identified in a range of neuropathologic conditions.** Spatial expression signatures of reactive glia populations (color scale), and concrete value distributions in the individual brain regions (right-side violin plots) are shown. Size of the gene list used to compute the expression signature is shown below, with the full gene lists attached inin Supp. Table 4. The expression signatures were quantified as difference in the average expression of this signature and a signature of iteratively randomly selected genes of similar expression levels (Methods).

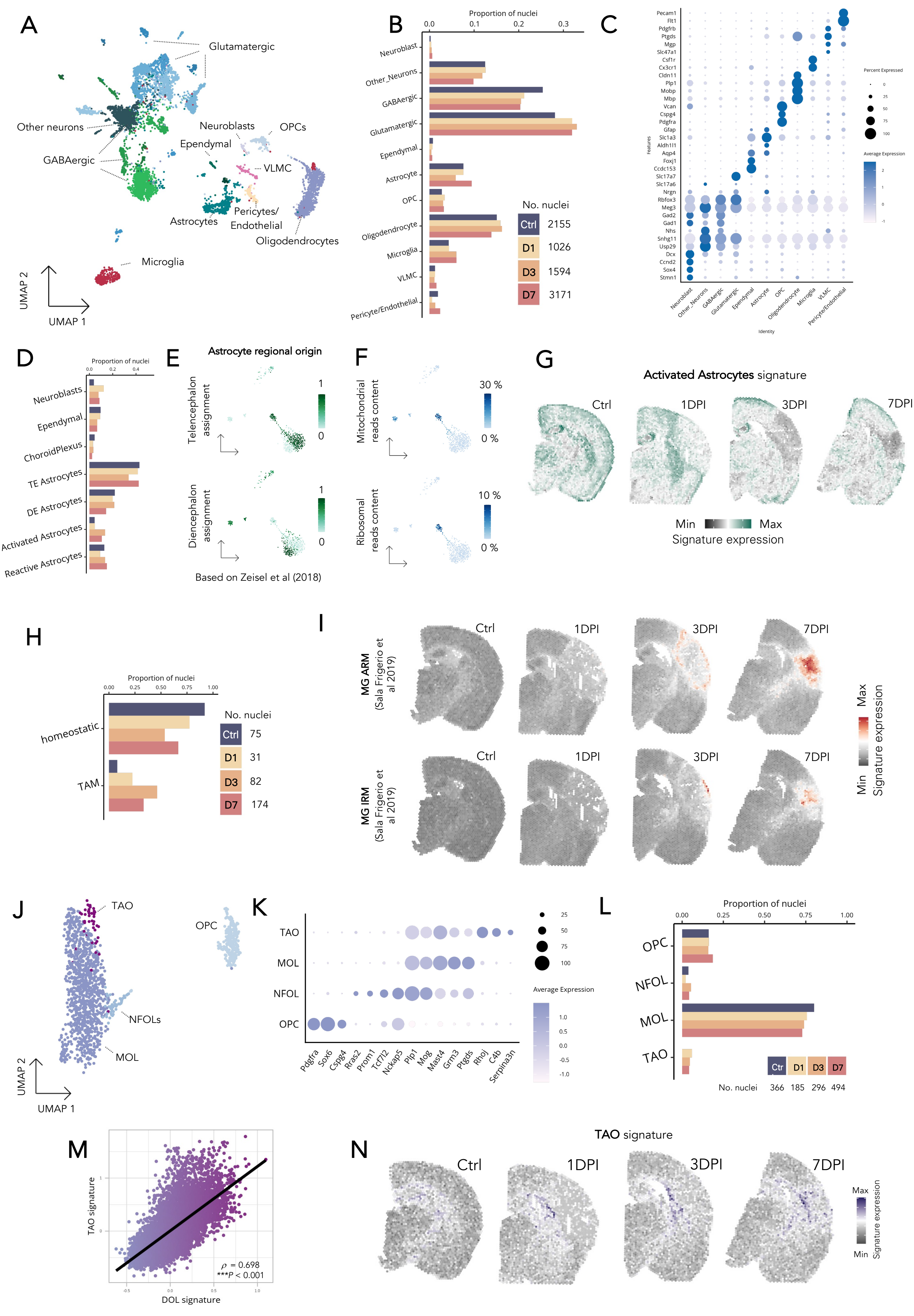

A
Glutamatergic
Other neurons
Neuroblasts
OPCs
Ependymal
GABAergic
VLMC
Pericytes/
Endothelial
Astrocytes
Oligodendrocytes
Microglia
UMAP 2
UMAP 1
B
Proportion of nuclei
0.0
0.1
0.2
0.3
Neuroblast
Other_Neurons
GABAergic
Glutamatergic
Ependymal
Astrocyte
OPC
Oligodendrocyte
Microglia
VLMC
Pericyte/Endothelial
No. nuclei
Ctrl 2155
D1 1026
D3 1594
D7 3171
C
Percent Expressed
Average Expression
Features
Identity
D
Proportion of nuclei
Neuroblasts
Ependymal
ChoroidPlexus
TE Astrocytes
DE Astrocytes
Activated Astrocytes
Reactive Astrocytes
E
Astrocyte regional origin
Telencephalon assignment
Diencephalon assignment
Based on Zeisel et al (2018)
F
Mitochondrial reads content
Ribosomal reads content
30 %
0 %
10 %
0 %
G
Activated Astrocytes signature
Ctrl
1DPI
3DPI
7DPI
Min
Max
Signature expression
H
Proportion of nuclei
homeostatic
TAM
No. nuclei
Ctrl 75
D1 31
D3 82
D7 174
I
MG ARM (Sala Frigerio et al 2019)
MG IRM (Sala Frigerio et al 2019)
Ctrl
1DPI
3DPI
7DPI
Max
Min
Signature expression
J
TAO
OPC
NFOLs
MOL
UMAP 2
UMAP 1
K
TAO
MOL
NFOL
OPC
Pdgfra
Sox6
Cspg4
Rras2
Prom1
Tcf7l2
Nckap5
Plp1
Mog
Mast4
Grm3
Ptgds
Rhoj
C4b
Serpina3n
Average Expression
L
Proportion of nuclei
OPC
NFOL
MOL
TAO
Ctr D1 D3 D7
No. nuclei 366 185 296 494
M
TAO signature
DOL signature
ρ = 0.698
***P < 0.001
N
TAO signature
Ctrl
1DPI
3DPI
7DPI
Max
Min
Signature expression

Supp. Figure 9 (previous page) – **Single-nucleus transcriptomic profiling of post-ischemic brain reveals rise of reactive glia populations.** A) UMAP of cell populations from post-MCAO mouse brains (n = 7946). B) Proportions of cell populations in the individual time points. C) Marker genes of cell populations. Expression level (color scale) and the percentage of the population expressing the marker (dot size) are shown. D) Proportions of astroependymal populations in the individual time points. E) ) Projection of the brain region origin. Mouse brain cell atlas of Zeisel et al (2018) was used for the label transfer. F) Percentage of mitochondrial (upper) and ribosomal (bottom) reads per cell. G) Spatial activated astrocytes signature is shown, calculated using the top 15 population marker genes (fold-change sorted quantified as difference in the average expression of this signature and a signature of iteratively randomly selected genes of similar expression levels (Methods). H) Proportions of microglial populations across the sampled timepoints. I) Spatial signatures of reactive microglial populations identified by Sala Frigerio et al. (2019), characterized by 33 genes (ARM) and 41genes (IRM), respectively. Full gene lists are in Supp. Table 4. J) UMAP of oligodendrocyte lineage cells from post-MCAO mouse brains (n = 1341 nuclei). Abbreviations: oligodendrocyte precursor cells, OPC; newly formed OLs, NFOL; mature OLs, MOL; trauma-associated OLs (TAO). K) Marker genes of cell populations. Expression level (color scale) across oligodendrocyte-lineage populations and the percentage of the population's cells expressing the marker (dot size) are shown. L) Proportions of astroependymal cell populations across the sampled timepoints. M) Correlation of increase between the TAO and disease-associated oligodendrocyte (DOL, Kenigsbuch et al., 2022) signatures in the spatial dataset, quantified with Spearman's rank correlation coefficient $\rho$. N) Spatial TAO signature is shown, calculated using the top 15 population marker genes (fold-change sorted), quantified as difference in the average expression of this signature and a signature of iteratively randomly selected genes of similar expression levels (Methods).

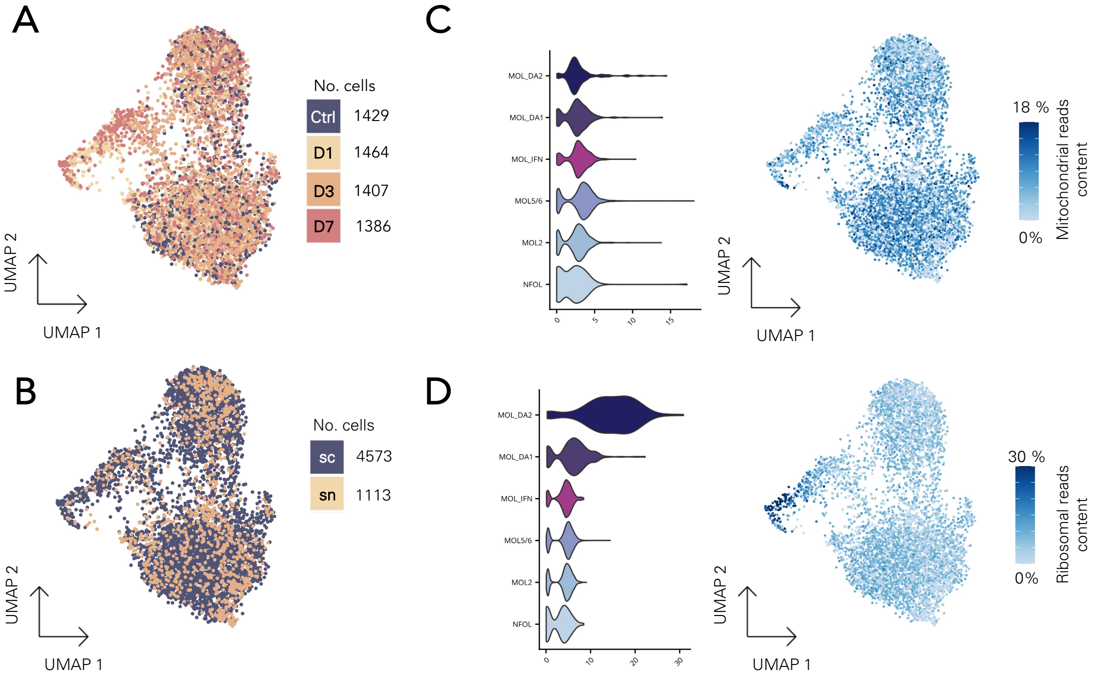


Supp. Figure 10 – **Integrative analysis of single-nucleus and single-cell post-MCAO oligodendrocyte datasets.** A) UMAP color-labeled based on the sampled timepoints. B) UMAP color-labeled based on the RNA-Seq method. single-cell, sc; single-nucleus, sn. C) Proportion of mitochondrial read content across the oligodendrocyte populations, on left the concrete values grouped by populations. D) Proportion of ribosomal read content across the oligodendrocyte populations, on left the concrete values grouped by populations.
